# Phylogenomics of monitor lizards and the role of competition in dictating body size disparity

**DOI:** 10.1101/2020.02.02.931188

**Authors:** Ian G. Brennan, Alan R. Lemmon, Emily Moriarty Lemmon, Daniel M. Portik, Valter Weijola, Luke Welton, Stephen C. Donnellan, J.Scott Keogh

## Abstract

Organismal interactions drive the accumulation of diversity by influencing species ranges, morphology, and behavior. Interactions vary from agonistic to cooperative and should result in predictable patterns in trait and range evolution. However, despite a conceptual understanding of these processes, they have been difficult to model, particularly on macroevolutionary timescales and across broad geographic spaces. Here we investigate the influence of biotic interactions on trait evolution and community assembly in monitor lizards (*Varanus*). Monitors are an iconic radiation with a cosmopolitan distribution and the greatest size disparity of any living terrestrial vertebrate genus. Between the colossal Komodo dragon *Varanus komodoensis* and the smallest Australian dwarf goannas, *Varanus* length and mass vary by multiple orders of magnitude. To test the hypothesis that size variation in this genus was driven by character displacement, we extended existing phylogenetic comparative methods which consider lineage interactions to account for dynamic biogeographic history and apply these methods to Australian monitors and marsupial predators. We use a phylogenomic approach to estimate the relationships among living and extinct varaniform lizards, incorporating both exon-capture molecular and morphological datasets. Our results suggest that communities of Australian *Varanus* show high functional diversity as a result of continent-wide interspecific competition among monitors but not with faunivorous marsupials. We demonstrate that patterns of trait evolution resulting from character displacement on continental scales are recoverable from comparative data and highlight that these macroevolutionary patterns may develop in parallel across widely distributed sympatric groups.

## Introduction

Organismal interactions provide an important selective force for evolution (Darwin 1859). On macroevolutionary time scales, interspecific interactions help drive the accumulation and distribution of diversity (Benton 1987). Common antagonistic interactions (e.g. competition) are suggested to facilitate the assembly of communities by encouraging ecological, behavioral, and morphological differentiation through character displacement (Brown and Wilson 1956; Sepkoski Jr 1996). This process has been repeatedly identified in insular adaptive radiations like Darwin’s finches, Caribbean anoles, and Lake Victoria cichlids, where young clades have rapidly diverged into many available phenotypes, ecologies, and/or behavioral syndromes (Schluter et al. 1985; Losos 1990; Grant and Grant 2006). While insular systems are instructive, they account for only a fraction of earth’s biodiversity, and it has been much more difficult to quantify the influence of competition at continental scales (Drury et al. 2018b). We therefore know little about how competition among organisms may influence the evolution of traits and distribution of most of life on earth.

The most obvious axis for differentiation between organisms is absolute size (Peters and Peters 1986). In animals, body size is often used as a proxy for guild, and because it dramatically affects life-history traits and ecology, it is the most commonly used measurement in macroevolutionary studies (Wilson 1975). Among terrestrial vertebrates, monitor lizards *Varanus* exhibit the greatest variation in body size within a single genus (Pianka 1995). Extant monitors include island giants like the Komodo dragon *V. komodoensis* (up to 3 m long and 100 kg), and desert dwarves like the short-tailed goanna *V. brevicauda* (0.2 m and 0.016 kg), which vary by orders of magnitude. In fact, while size estimates vary, the recently extinct Australian monitor *Varanus (Megalania) priscus* may have dwarfed even the Komodo dragon, reaching lengths of over four meters (Wroe 2002; Conrad et al. 2012). Despite a conservative body plan, monitor lizards are ecologically diverse and can be found at home in trees, among rocks, in burrows, and swimming through watercourses and even the open ocean (Pianka 1995). Though there are roughly 80 described monitors, the greatest morphological diversity is concentrated in the 30 or so Australian species (Uetz and Hošek 2019). All Australian monitors are hypothesized to constitute a single radiation that likely dispersed from Sundaland into Sahul (Australopapua), though the timing and biogeographic history of this group remains uncertain (Vidal et al. 2012). Such incredible diversity in body size begs the question, what has driven it?

Over the years, researchers have suggested that this disparity is the result of habitat partitioning (Collar et al. 2011), or release from competition with carnivoran mammals (Pianka 1995; Sweet and Pianka 2007). However, no one has yet investigated whether variation in monitor body sizes is instead the result of character displacement through competition, either with other *Varanus* or other large carnivores with which they may vie for resources. This is likely due to the fact that probabilistic trait evolutionary models largely remain ignorant of such interactions even though they are ubiquitous (Harmon et al. 2019). Only recently have methods for modelling continuous traits attempted to take into account the influence of lineages on one another (Drury et al. 2016; Manceau et al. 2017; Adams and Nason 2018; Quintero and Landis 2019).

In Australia, monitor lizards are not the only radiation of terrestrial vertebrate predators. A similarly diverse co-distributed group are the carnivorous and omnivorous marsupial mammals. Dasyuromorphians and peramelemorphians cover a similar breadth in range and body size, inhabiting deserts and closed forests, ranging from the tiny *Ningaui* up to the recently extinct canine-convergent *Thylacine*. Outside of Australia, there is evidence to suggest varanid lizards may compete either directly (through predation) or indirectly (vying for resources) with small-to-moderate sized carnivo-rans, and this may explain the lack of small monitors west of Wallace’s Line (Sweet and Pianka 2007). This presents the question of whether or not Australian monitors and their marsupial neighbors have influenced the size evolution of one another, and if this signature may be discernible from comparative data.

In order to address these macroevolutionary questions on the origins and diversity of varanid lizards, it is essential to first construct a reliable time-scaled phylogeny. Relationships among *Varanus* have been reconstructed historically through a number of morphological and molecular methods, but recovered subgeneric relationships have been notoriously inconsistent (Fuller et al. 1998; Ast 2001; Fitch et al. 2006; Conrad et al. 2012; Vidal et al. 2012; Lin and Wiens 2017). We generated a nuclear exon capture dataset and combined it with existing morphological data to build a comprehensive phylogenetic hypothesis for *Varanus* in a combined evidence framework, incorporating both fossil and extant taxa. Our phylogenetic estimates are well-resolved at multiple taxonomic levels and we use them to reconstruct the global biogeographic history of varaniform lizards, then focus on the evolution of body size among Australian taxa. To address the influence of competition on size evolution, we extend a series of novel comparative phylogenetic models. These include models which integrate continental biogeographic history (not just contemporary distribution), and the possibility of competition with another group of highly diverse Australian carnivores.

## Materials & Methods

Walkthroughs of the data, code, analyses, and results are available in the *Supplementary Material*, on GitHub at www.github.com/IanGBrennan/MonitorPhylogenomics, and from the Dryad Digital Repository: http://dx.doi.org/10.5061/dryad.tx95x69t8

### Molecular Data Collection

We assembled an exon-capture dataset across 103 *Varanus* specimens representing 61 of 80 currently recognized species. This sampling covers all nine subgenera and major clades of *Varanus*, as well as recognized subspecies, and known divergent populations. We included four additional non-varanoid anguimorphs (*Elgaria*, *Heloderma*, *Shinisaurus*, *Xenosaurus*), a skink (*Plestiodon*), and tuatara (*Sphenodon*) as outgroups. Nuclear exons were targeted and sequenced using the Anchored Hybrid Enrichment approach (Lemmon et al. 2012), and resulted in 388 loci (average coverage 350 loci, min = 112, max = 373) totalling ∼600 kbp per sample (Fig.S7).

### Morphological Sampling

In addition to novel phylogenomic sampling, we included morphological data collected by Conrad et al. (2011). We chose to exclude a number of characters added to this matrix in Conrad et al. (2012) because of extensive missing data and uncertain homology. We filtered the data matrix using an allowance of 50% missing data per character, excluding characters above this threshold, and removed taxa with greater than 70% missing data, as we found these samples to be disruptive in exploratory analyses. We removed invariant characters from the remaining data to conform to assumptions of the MKv model, resulting in a final morphological matrix comprising 303 characters. Disruptive samples—often called ‘rogues’—are not limited to those with large amounts of missing data. To identify if rogue taxa are causing topological imbalances in our phylogenetic hypotheses, we applied RogueNaRok (Aberer et al. 2012) to initial combined evidence analyses, identified rogues, and removed them for downstream analyses. Morphological sampling includes 55 extant *Varanus*, as well as the extinct *V. priscus*. A number of extant and fossil outgroups are included to sample the closely related groups Helodermatidae (*Heloderma suspectum*), Lanthanotidae (*Lanthanotus borneensis*, *Cherminotus longifrons*), Paleovaranidae (formerly Necrosauridae) (*Paleovaranus (Necrosaurus) cayluxi*, *P. giganteus*, *‘Saniwa’ feisti*) (Georgalis 2017), Shinisauridae (*Shinisaurus crocodilurus*), and uncertain varaniform lizards (*Aiolosaurus oriens*, *Ovoo gurvel*, *Telmasaurus grangeri*, *Saniwides mongoliensis*).

### Phylogenetic Analyses

We reconstructed a partitioned concatenated species tree and individual genealogies for our exon capture data (n=388) under maximum-likelihood in IQTREE (Schmidt et al. 2014), allowing the program to assign the best fitting model of molecular evolution using PartitionFinder, then perform 1,000 ultrafast bootstraps (Haeseler et al. 2013). We then estimated the species tree using the shortcut coalescent method ASTRAL III (Zhang et al. 2017), with IQTREE gene trees as input. Further, we also estimated species trees using the full multispecies coalescent (MSC) and fossilized birth-death MSC (FBD-MSC) models implemented in StarBEAST2 (Ogilvie et al. 2016). Computational limitations under the MSC required we reduce the input data size and so we summarized per-locus informativeness using AMAS (Borowiec 2016), then used custom scripts to sort the loci sequentially by (*i*) missing taxa per alignment, (*ii*) number of variable sites, and (*iii*) AT content. We then chose the first three sets of twenty loci (1–20; 21–40; 41–60) as representatives of the most informative and complete loci, and used them to build our phylogeny (Fig.S8).

Phylogenetic reconstruction under the FBD-MSC allowed us to jointly infer a molecular and morphological species tree, and divergence times using structured node and tip date priors (Supplementary Material: “Node Priors and *Varanus* in the Fossil Record”; Table S8). Morphological data were modelled under the Mkv model, a special case of the Mk model (Lewis 2001)—the most commonly used model for discrete morphological data. We partitioned morphological characters by differing numbers of states following Gavryushkina et al. (2017). All StarBEAST2 analyses were run for four independent chains under uncorrelated relaxed lognormal (UCLN) and strict molecular clocks for 1 billion generations and sampled each 5×10^5^ generations, to assess convergence among runs. To further inspect our prior assumptions we ran all analyses under the priors only and compared against empirical runs. We inspected the MCMC chains for stationarity (ESS > 200) using Tracer v1.7.0 (Rambaut et al. 2018), and discarded the first 10-40% of each run as burn-in as necessary before combining runs. Combined evidence analyses may be biased by difficulties in accurately modelling morphological evolution (Puttick et al. 2017; Luo et al. 2018; Goloboff et al. 2018). In contrast to molecular sites or loci, morphological characters are likely more often correlated (Billet and Bardin 2018), nonhomologous (Baum and Donoghue 2002), or evolving under dramatically different mechanisms (Goloboff et al. 2018), and may disrupt our best efforts at reconstructing phylogeny, divergence times, and rates of evolution. To address this, we also estimated divergence dates using an “extant-only” approach, limiting the sampling to living taxa with molecular data, and used the multispecies coalescent model implemented in StarBEAST2, using the same clock and substitution models, and chain lengths as above.

Fossil taxa are almost always assumed to represent terminal tips that have since gone extinct. To test this assumption, we allowed fossil taxa to be identified as terminal or stem lineages using the *Sampled Ancestors* package implemented in StarBEAST2. Using our prior-only analyses we calculated Bayes factors (BF) for each fossil taxon to test competing hypotheses (ancestor or tip). We used a threshold of log(BF) > 1 to identify sampled ancestors, log(BF) < −1 to recognize terminal taxa, and −1 < log(BF) < 1 taxa were categorized as equivocal.

### Biogeographic History

*Varanus* lizards have been variously hypothesized to have originated in Asia (Keast 1971; Estes 1983; Fuller et al. 1998; Jennings and Pianka 2004; Amer and Kumazawa 2008; Vidal et al. 2012; Conrad et al. 2012), Africa (Holmes et al. 2010), or Gondwana (Schulte et al. 2003) with conclusions largely based on which taxa were included, and the timing of varanid divergence events. We used *BioGeoBEARS* (Matzke 2014) to infer the biogeographic history of varanids and kin, dividing their range into seven major regions: North America, Europe, Sundaland/Wallacea, AustraloPapua, Africa/Arabia, West Asia (Indian subcontinent and surrounds), and East Asia (China, Mongolia, mainland Southeast Asia). As input we used the maximum clade credibility tree from our combined evidence analyses. Because of the deep evolutionary history of this group we took plate tectonic history into account by correcting dispersal probability as a function of distance between areas. We estimated distances between areas and continents through time at five million year intervals from 0–40 million years ago, then ten million year intervals from 40–100 million years, using latitude and longitude positions from GPlates (Boyden et al. 2011), and calculated pairwise distance matrices using the R package *geosphere* (Hijmans 2016). Additionally, we limited the model-space by providing information about area adjacency. For each time period, we removed unrealistic combinations of ranges (e.g. North America + AustraloPapua), with the aim of recovering more realistic biogeographic scenarios.

To understand the spatial evolution of *Varanus* in Australia, we used a Bayesian method *rase* (Quintero et al. 2015) which assumes a Brownian motion diffusion process to infer ancestral ranges as point data. We downloaded occurrence records for all continental Australian *Varanus* species from the Atlas of Living Australia (ala.org), curating the data for erroneous records, then trimmed our input tree down to just Australian taxa. We ran *rase* for 10,000 generations, sampling each 10th generation, then discarded the first 10% (100 samples) as burn-in, leaving 900 samples. We inspected the traces of the MCMC chains for stationarity using *coda* (Plummer et al. 2006).

### Signature of Character Displacement

Ecological communities are generally thought to assemble under opposing processes of habitat filtering and interlineage competition. Filtering is suggested to select for species with similar phenotypes, resulting in conservatism or convergence, whereas competition is expected to result in greater phenotypic disparity. These expectations can be tested by investigating the functional diversity of communities across the landscape. We divided the Australian continent into half-degree cells, and created a site by species matrix using the ALA distribution data for (**i**) monitor lizards and again for (**ii**) monitors and dasyuromorph/peramelemorph marsupials together. We estimated the functional diversity for the two data sets using the package *FD* (Laliberté et al. 2014) and Rao’s Quadratic, using body size as the trait of interest. We then estimated functional diversity for each inhabited cell 100 times using a dispersal null metric model which sampled from nearby cells assuming a probability proportional to the inverse of the distance from the focal cell. To compare observed and simulated functional diversities, we calculated standardized effect sizes (SES) for each cell, and a mean SES across the continent with 95% confidence intervals.

### Modelling Body Size Evolution with Competition

Only within the past few years have phylogenetic comparative methods (PCMs) begun to account for the interaction of lineages on trait evolution. Conceptual work by Nuismer and Harmon (2015) led to the development of the *Matching Competition* (*MC*) model by Drury et al. (2016), which infers an interaction parameter (*S*) dictating attraction towards or repulsion from the mean trait value of interacting lineages. This was extended by Drury et al. (2018b) to incorporate interactions matrices which limited interactions to only codistributed species. We build upon this framework by expanding the biogeographic information to include temporally and spatially dynamic ranges for ancestral taxa (inferred from *rase*, example in Fig.S6). In natural ecosystems, many different organisms compete for the same resources, so accounting for competition only within a single group is perhaps unrealistic. To address this issue, we consider the influence of another broadly distributed group of like-sized carnivores and omnivores, dasyuromorphian and peramelemorphian marsupials, on the size evolution of Australian monitor lizards. To test this hypothesis we begin by trimming the marsupial phylogeny of Brennan and Keogh (2018) down to just the faunivorous clades, from which we also dropped *Myrmecobius* because of its unusual ecology. We collected body size (mm) information for marsupials from Pantheria (Jones et al. 2009), and monitors from the literature (Wilson and Swan 2013). Manceau et al. (2017) introduced a framework for estimating the effect of one clade on the trait evolution of another, incorporating two phylogenetic trees, referred to as the *Generalist Matching Mutualism* (*GMM*) model. This is essentially a two-clade extension of the *MC* model, which makes the assumption that the evolution of trait values in clade A are the result of interactions *only* with lineages in clade B, and vice versa. The *GMM* model however makes two very basic assumptions that we expect do not fit our data: (**1**) interactions between phenotypes are limited to interclade (between trees) matching or competition, meaning there is no influence of intraclade (within tree) interactions, and (**2**) that all contemporaneous lineages are interacting, regardless of geographic distribution. To address these assumptions, we developed and fit a series of models that expand on the interaction parameter *S* and incorporate biogeography to provide more realistic models of trait evolution. We present summaries and graphical descriptions of these models in Fig.4 and the *Supplementary Material* (Fig.S1). Further, we test if size evolution is instead dictated by non-ecological processes, by employing standard models of trait evolution, Brownian Motion **BM** and Ornstein Uhlenbeck **OU**. Using these traditional null models, we can again ask if monitor and dasyuromorphian size has evolved under similar or independent rates using *ratebytree* in *phytools*, though we also provide implementations of shared BM and OU models in the *RPANDA* framework—**CoBM** and **CoOU**. To compare against an alternative hypothesis of varanid size evolution (Collar et al. 2011) where variation is dictated by habitat use, we also fit a multi-optima (OUM) model in *OUwie* (Beaulieu et al. 2012).

To incorporate historical and contemporary biogeography, we extended our *rase* analyses to marsupials with data collected from the ALA. We designed a number of custom scripts and functions to process the spatial data and model objects including extensions of the ‘CreateGeoObject’ of *RPANDA*. Our functions ‘CreateGeoObject_SP’ and ‘CreateCoEvoGeoObject_SP’ produce *RPANDA* GeoObjects that take as input a tree, spatial distribution data in latitude/longitude format, and a post-processed *rase* object. Internally, these functions use the packages *sp* and *rgeos* to translate spatial data into spatial polygons representative of species distributions. Then, at each cladogenetic event, we determine the pairwise overlap of all contemporaneous lineages to construct our GeoObject (see Fig.S6). The ‘CreateCoEvoGeoObject_SP’ function has adapted this process for two trees, to be applied to *GMM* -type models.

### Model Behavior and Identifiability

The ability to identify competition and estimate associated parameters using process-based models has been tested extensively previously (Drury et al. 2016, 2018a, 2018b). From this we know that the ability to recover competitive models and estimate the interaction parameter *S* —when it is the generating process—is strongly linked to the absolute value of *S*, and to a lesser degree the size of the phylogeny. Parameter estimate and recovery of *S* can also be highly influenced by the incorporation of stabilizing selection (*ψ* or *α*), with the two parameters working agonistically in instances of competition (-*S*), and synergistically in mutualistic circumstances (+*S*). To ensure that we can accurately identify our models and estimate parameter values, we undertook a focused simulation exercise. Following the advice of Manceau et al. (2017), we simulated data directly onto our Australian monitor and marsupial trees under the same models we fit to our empirical data: BM_shared_, OU_shared_, CoEvo, CoEvo_all_, CoEvo_split_, JointPM_geo_, and CoPM_geo_. We used the *RPANDA* function ‘simulateTipData’ to simulate body size data under all specified models, keeping the empirical biogeography constant. Specifics of the generating parameter values are noted in the Table S3. We then iteratively fit the models to our simulated data, and compared fit using AICc and plotted AICc weights. To determine the ability to accurately recover parameter values, we then compared estimated to simulated values under each model.

## Results

### Phylogenetics of Monitor Lizards and Kin

Topologies estimated across maximum-likelihood (IQTREE Schmidt et al. (2014)), shortcut-coalescent (ASTRAL Zhang et al. (2017)), and Bayesian multispecies coalescent (StarBEAST2 Ogilvie et al. (2016)) methods are highly concordant and generally strongly supported (Fig.S3,1). Contentious nodes are limited to some subspecific (*Varanus gouldii*, *V. panoptes*) and interspecific relationships (*V. salvator* complex) which occur across a number of extremely short branches with low gene concordance factors, indicating both low information content and confidence. All analyses support the monophyly of *Varanus* and anguimorphs, and unite the Shinisauridae with the Helodermatidae, Anguidae, and Xenosauridae along a short internal branch. The Varanidae is sister to this group.

Interestingly, much of our trees are consistent with the first molecular phylogenies of *Varanus* proposed by Fuller et al. Fuller et al. (1998) and Ast et al. Ast (2001) two decades ago. Our results verify the monophyly of African and Arabian monitor lizards, and contrary to other recent studies (Lin and Wiens 2017), support the monophyly of both *Psammosaurus* and *Polydaedalus* subgenera. Our data support a geographically widespread clade comprising Philippines (*Philippinosaurus*) and tree (*Hapturosaurus*) and mangrove monitors (*Euprepiosaurus*), with water monitors (*Soterosaurus*) and species from the Indian subcontinent (*Empagusia*). We return a well resolved clade of Indo-Australopapuan monitors comprising the crocodile monitor (*Papuasaurus*), and the subgenera *Varanus* and *Odatria* (the dwarf monitors). Further, we record the first phylogenetic placement of the engimatic monitor *V. spinulosus* (*Solomonsaurus*) as sister to the Asian and Pacific clade, and confidently place *V. gleboplama* as sister to the rest of *Odatria*.

Dating estimates from our combined evidence and node-calibrated molecular analyses in Star-BEAST2 agree on the timing of *Varanus* divergences. They suggest an origin of varanids (split between Varanidae and Lanthanotidae) in the mid-to-late Cretaceous (80–100 ma), and an early-to-mid Oligocene (28–35 ma) origin for the crown divergence of extant *Varanus*. These dates are comparable with recent estimates from the literature (Lin and Wiens 2017; Pyron 2017), and younger than previous estimates (Vidal et al. 2012; Portik and Papenfuss 2012) which used stem varanids to calibrate the crown (Fig.S4). Ten fossil taxa form relatively poorly resolved higher-order relationships, with the Palaeovaranidae (formerly Necrosauridae) forming a clade with the Lanthanotidae (*Lanthanotus*, *Cherminotus*), together as sister to the Varanidae (*Varanus*, *Saniwa*). *Varanus priscus*, which is generally considered an extinct relative of the Indo-Australopapuan clade of giant monitors including *V. varius*, *V. komodoensis*, and *V. salvadorii*, is consistently placed in the Australian radiation. Given the existing morphological data, the majority of fossil taxa are recovered as tips in our analyses (Fig.S10).

### Biogeography and Community Assembly

Global biogeographic analysis of *Varanus* and allies suggests an origin of varaniform lizards in East Asia, with dispersals west across Laurasia into Europe, and east into North America. The origin of the genus *Varanus* is equivocal (Fig.S11), but likely followed a similar pattern, with independent clades dispersing west through the Middle East and into Africa and Europe, and south and east through Southeast Asia, Sundaland, and into Indo-Australia. After reaching the western and eastern extents of their range, both the African and Australopapuan clades appear to have begun dispersals back towards their origins. This has resulted in *V. yemenensis* extending across the Red Sea into the Arabian Peninsula, and *V. komodoensis* and members of the *V. scalaris* complex reaching back into Wallacea. A DEC model incorporating dispersal probability as a function of distance is strongly preferred (AIC = 170.66, *x* = −0.682) over the traditional DEC model (AIC = 186.04, ΔAIC = 15.38).

Biogeographic reconstruction of Australian *Varanus* reveals an origin spread across much of northern and central Australia (Fig.3). Considering northern Australia was the most likely colonization point for monitors, it makes sense that our analyses of community structure highlight this area as the center of greatest species richness for *Varanus*, with up to eleven species recorded in some half-degree grid cells. Taken together with dasyuromorph and peramelemorph marsupials, we again see high richness in the Top End, but also note species richness hotspots in the Central Deserts and the Pilbara regions. These regions are functionally diverse for monitors as well, but much less so for communities of marsupials and monitors analyzed jointly. Overall, we find support for overdispersion in trait values in the monitor-only dataset. Across Australia functional diversity of most communities is greater than expected under our null model (mean SES across all cells for monitors = 0.07 *±* 0.05;). Functional diversity is greatest in monitor communities of moderate-to-high (3–7 spp.) richness (mean SES = 0.45 *±* 0.13), and lower than we would expect under our null model in communities of only two species (mean SES = −0.16 *±* 0.08) (Table S9). In contrast, communities of monitors and marsupials together have estimates of functional diversity consistently lower than expected under the null model (mean SES across all cells = 1.2 *±* 0.26) (Table S10).

### Modelling Body Size Evolution

We extend a coevolutionary comparative method framework (Manceau et al. 2017) to incorporate historical biogeography and estimate the influence of lineage interactions on trait evolution. Comparison of traditional models of trait evolution (Brownian Motion, Ornstein Uhlenbeck) with those that incorporate interactions among lineages decisively favors interactive models (AICc weight 94%) (Figs.4, S12). These models can be broadly divided into those which estimate the interaction parameter *S* from occurrences (**1**) within clades (*S* _intra_), (**2**) between clades (*S* _inter_), or (**3**) both. We find greatest support for models that estimate interactions only within clades (Fig.4). Support for the best-fitting model *CoPM_geo_*—which fits only a single *S* _intra_ parameter for *both* trees—suggests that the strength of intraclade interactions can not be differentiated between the two groups. Across fitted models that estimate *S* _intra_, we inferred negative values of *S*, supporting competitive interactions in both monitors and marsupials, *S* _intra_ = −0.043 *±* 0.005.

Support for the *CoPM_geo_* model also comes indirectly from parameter estimates of the *CoEvo_split_* model. In fitting the *CoEvo_split_* model, which estimates separate inter- and intraclade interaction parameters (*S_inter_*, *S_intra_*), we estimate a weak positive *S_inter_* parameter of 0.0043. This parameter estimate is small enough to likely be biologically meaningless, and with *S_inter_ ≈* 0 the *CoEvo_split_* model collapses to *CoPM_geo_* (see *Supplementary Material—Nested Models*). This suggests that interclade interactions between marsupials and varanids are indistinguishable from these data.

Results of our model identifiability exercise indicate that all proposed models can be recovered under realistic circumstances (Fig.S13). Because a number of these are nested forms of one variety or another, when simulated values of *S* (as *S_1_* or *S_2_*) approach 0, some models may be incorrectly conflated. Consistent with previous assessment (Drury et al. 2016), we also find that the accuracy of estimated *S* is directly related to the absolute value of *S*, with greater values of *S* being more precisely recovered (Fig.S14).

## Discussion

Competitive interactions are expected to impact diversity by influencing species ranges, and influence phenotypic and behavioral evolution through character displacement (Brown and Wilson 1956; Benton 1987). *Varanus* represent a diverse group of lizards with exceptional variation in body size and ecologies (Fig.2). To investigate the role of competition in size evolution in monitors, we started by building a phylogenomic hypothesis of living and extinct varanids and their allies. By using a total evidence dating approach we were able to take advantage of both molecular and morphological data to incorporate fossil taxa, and reconstruct the global biogeography of varaniform lizards. Focusing on the Australian continent, we used a temporally dynamic Brownian Motion dispersal process to infer ancestral ranges for monitor lizards and co-occurring marsupial predators. We then quantified the functional diversity of monitor communities, and monitor–marsupial communities to address how these assemblages are structured. Finally, we developed and implemented a number of comparative models to account for interspecific interactions and estimate competition among monitors *and* with dasyuromorphian marsupials. Results of our comparative modelling provide a compelling case for considering competition in phylogenetic comparative methods (PCMs) of trait evolution.

**Figure 2:**
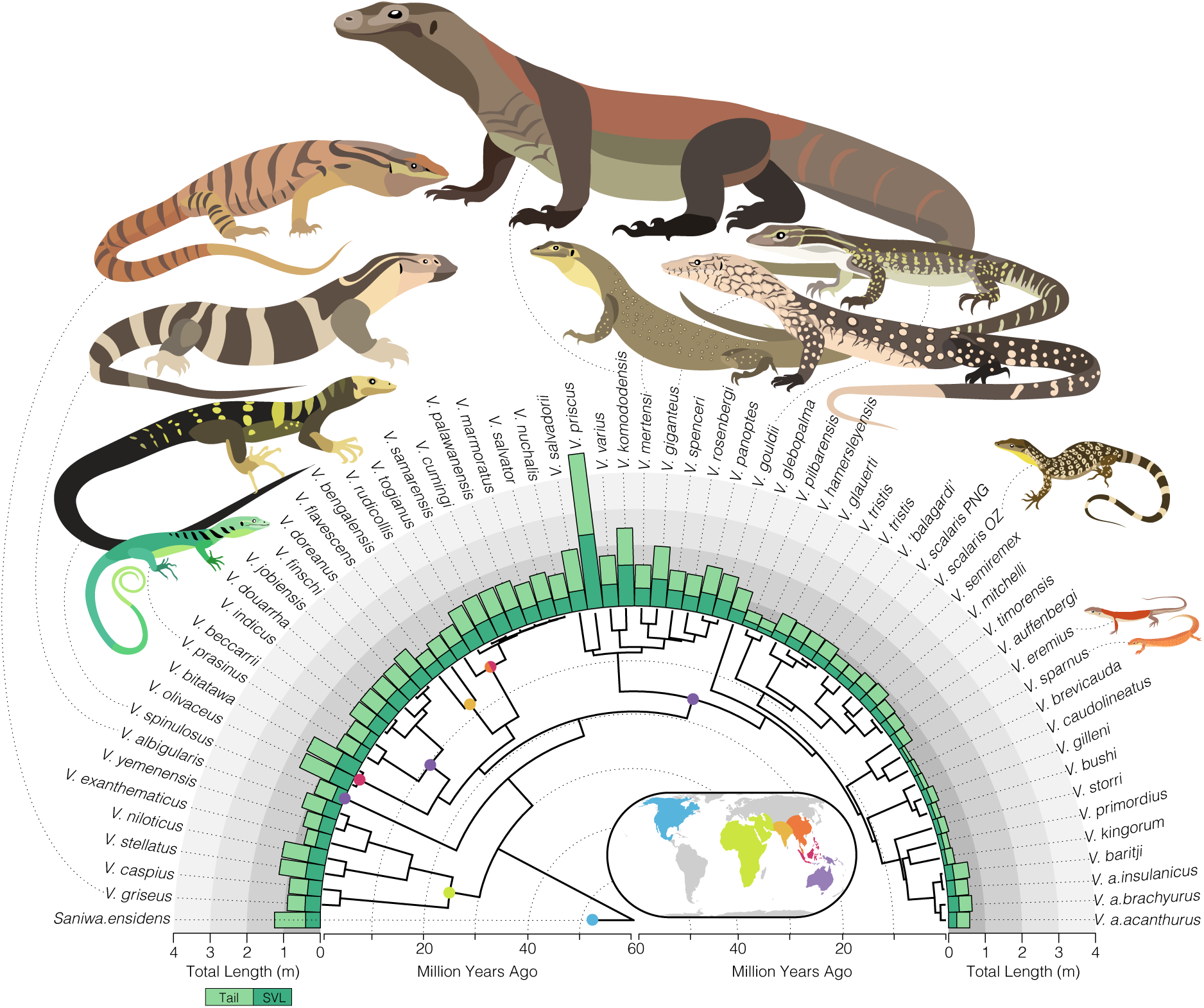
Body size among *Varanus* species varies across multiple orders of magnitude. Bar plots at tips of the tree show total length of sampled monitor lizards broken down into snout-vent length (SVL) and tail length. The smallest monitor species *Varanus sparnus* reaches just over 200 mm long from snout to tail tip and may weigh only 20 g, while the largest living species *Varanus komodoensis* can reach well over 2 meters long (2000+ mm) and top the scales at 100 kg (100,000 g). By all accounts, the recently extinct *Varanus priscus* was even larger than the Komodo dragon and may have reached over 4 m long (Wroe 2002; Conrad et al. 2012). Inset map shows a rough global distribution of monitor lizards and the extinct relative *Saniwa ensidens*. Colored circles at nodes indicate primary distribution of the major clades of *Varanus* and correspond to distributions on the map (blue–North America; green–Africa and the Middle East; light orange–Indian Subcontinent; dark orange–Indochina and China; red–Sundaland and Wallacea; purple–AustraloPapua).

**Figure 3:**
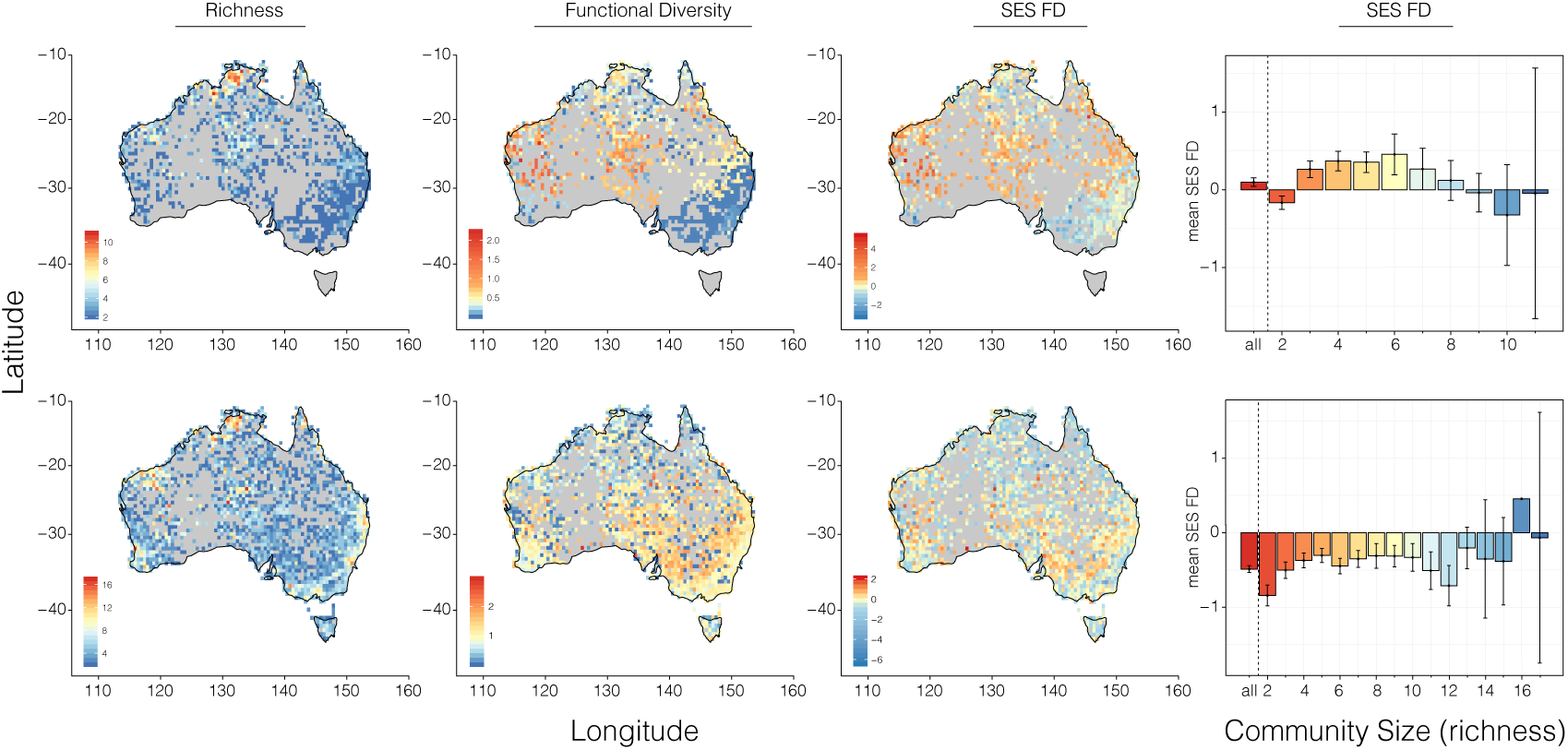
Maps of Australia showing patterns of richness (number of species) and functional diversity for monitor lizards (top row) and for monitor lizards and faunivorous marsupials together (bottomr row). Values were calculated and plotted for half-degree squares, with warmer colors indicating greater values—but note different scales for each plot. The left plots display species richness across the landscape and center-left plots show absolute values for functional diversity (FD—Rao’s Q). Center-right plots show the standardized effect size (SES) of functional diversity when compared to the dispersal-corrected null model, and right plots show how the mean standardized effect sizes vary across communities of varying richness. In communities of moderate richness (3–7 spp), functional diversity is overdispersed in monitor lizards, suggesting character displacement. Functional diversity is almost always underdispersed when considering monitors and marsupials in communities together.

**Figure 4:**
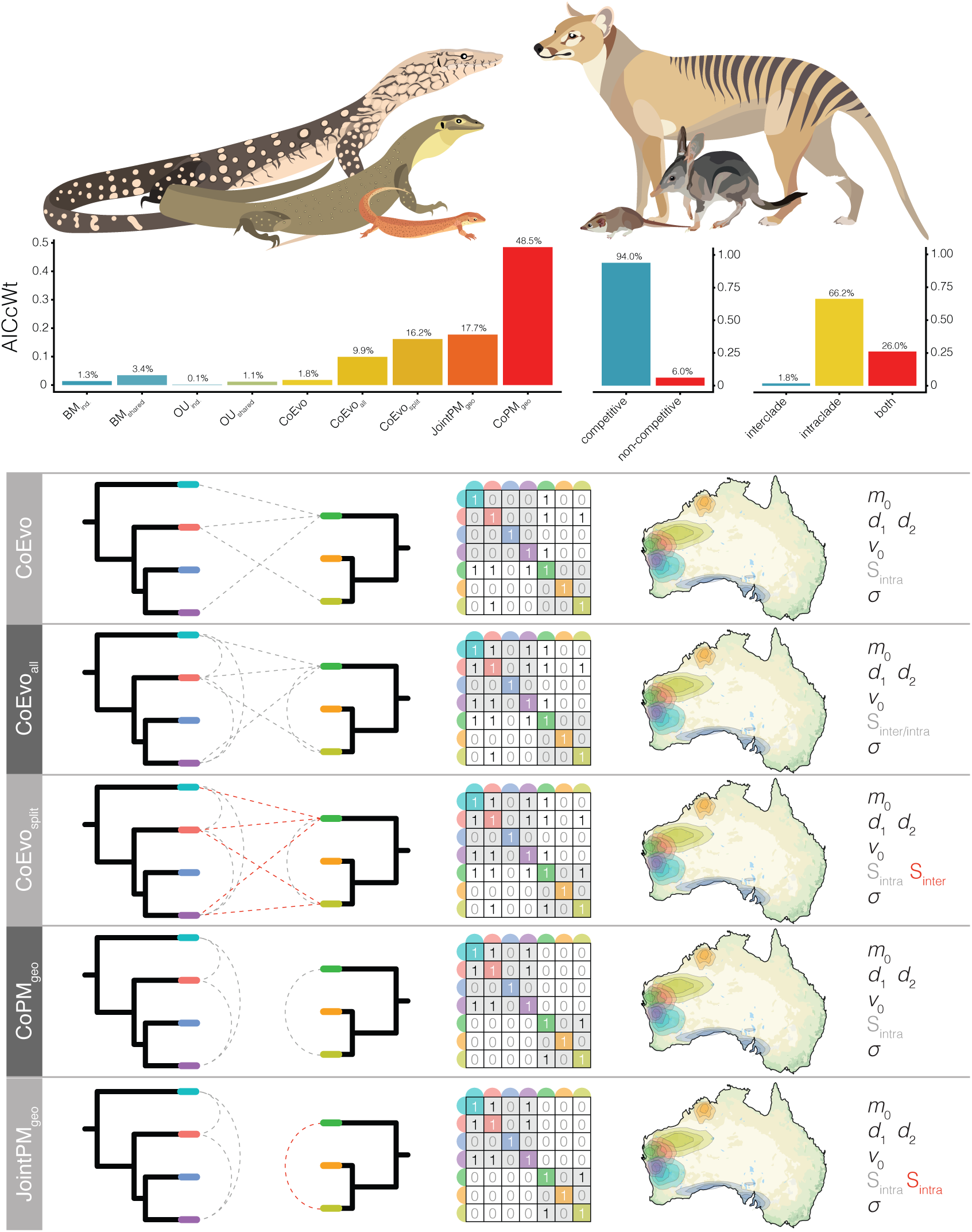
Comparative model fitting highlights the importance of incorporating interactions when modelling body size evolution of monitor lizards and faunivorous marsupials. Top, examples of Australian monitor lizards (*Varanus giganteus*, *V. mertensi*, *V. sparnus*) and marsupials (*Thylacinus cynocephalus*, *Macrotis lagotis*, *Pseudantechinus bilarni*), drawn roughly to scale. Middle, modelling competition vastly improves model fit, but size evolution appears largely driven by intraclade evolution and not competition between monitors and mammals. Bottom, hypothetical schematic components of biogeographically-informed lineage-interaction comparative models for two clades. Each model is named at left, followed by a diagram of the the two trees with interlineage interactions allowed under the given model designated by dashed lines. If more than one interaction parameter *S* is estimated, it is denoted by red dashed lines. The contemporary summary of these interactions are presented in the interaction matrix *P*, and the estimated parameters are listed at far right. Maps show the distribution of the taxa used in these examples, and inform the interaction matrices.

### Phylogenetic Relationships and Origins

Relationships between anguimorph lizard groups have been contentious, particularly with regard to the placement of fossil taxa (Conrad 2008; Conrad et al. 2011; Pyron 2017). Different datasets have supported strongly competing hypotheses including a monophyletic Varanoidea (Varanidae, Shinisauridae, Monstersauria) (Gauthier et al. 2012), paraphyly of Varanoidea with regards to Anguidae, and even sister relationships between Varanidae and Mosasauria (Conrad 2008) or Varanidae and Serpentes (Hejnol et al. 2018). Existing hypotheses about relationships among these groups appear highly sensitive to the data used, with conflicting molecular and morphological signals (Pyron 2017; Hejnol et al. 2018), and even incongruences between different morphological datasets (Conrad 2008; Conrad et al. 2011; Gauthier et al. 2012; Pyron 2017). Much of this likely has to do with the fragmentary nature of many fossil taxa, morphological models of character evolution, and previous reliance on mitochondrial DNA of extant taxa. Our reanalysis of these morphological data in concert with novel phylogenomic data are largely consistent with previous assessments, however we provide new insights into the phylogenetics of living members of *Varanus*.

One of the most intriguing results from our data is the the phylogenetic placement of *V. spinulosus* Although it is not wholly unexpected (Ziegler et al. 2007b, 2007a; Bucklitsch et al. 2016), it is not affiliated with the subgenus *Varanus* (Sweet and Pianka 2007) or with *Euprepiosaurus* (Harvey and Barker 1998). Instead, we place *V. spinulosus* alone on a long branch between the African and Asian monitors, and corroborate the previous erection of a unique subgenus *Solomonsaurus* (Bucklitsch et al. 2016). The phylogenetic position of *V. spinulosus* is remarkable given that it is a Solomon Islands endemic, meaning it likely made a considerable over-water dispersal or island hopped to the Solomons only shortly after their formation ∼30 Ma (Hall 2002). This corroborates the intriguing observation that relatively young Melanesian islands have long been sources for ancient endemic diversity (Pulvers and Colgan 2007; Heads 2010; Oliver et al. 2017, 2018). It also suggests at least three independent dispersals of *Varanus* across Wallace’s line, and a convoluted history of movement throughout the Indo-Australian region.

Our phylogeny of *Varanus* also highlights the adaptive capacity of these amazing lizards (Fig.2, S5). For example, the perentie *V. giganteus* is the largest extant Australian lizard, reaching well over two meters long, while remaining extremely thin. Its sister species *V. mertensi* in contrast, is a heavy bodied semiaquatic lizard built for the watercourses of northern Australia. Together, these species are sister to a group of sturdy terrestrial wanderers—the sand goannas—*V. gouldii*, *V. panoptes*, *V. rosenbergi*, and *V. spenceri*. In roughly five million years, these monitors diverged broadly both ecologically and morphologically, and spread across Australia’s landscape. In the process of diversifying, monitor lizards have also converged repeatedly on ecological niches and body plans. There are at least four different origins of amphibious monitors (*V. salvator*, *V. mertensi*, *V. mitchelli*, *V. niloticus* groups), and four or more origins of arboreal species (*V. prasinus*, *V. gilleni*, *V. salvadorii*, *V. olivaceous*, *V. dumerilii* groups), emphasizing the ability of monitors to fill available niches.

A number of phylogenetic questions evade our sampling, and largely concern the population genetics of known species complexes. These include the *V. acanthurus*, *V. doreanus*, *V. griseus*, *V. indicus*, *V. jobiensis*, *V. prasinus*, *V. salvator*, *V. scalaris*, and *V. tristis* groups, of which most have recognized subspecies, very closely related species, or are paraphyletic in our data (Fig.1). Some of these taxa have experienced dramatic taxonomic growth in recent years as a result of more extensive sampling, and are sure to present exciting phylogeographic and systematic stories when the right data and sampling are paired together.

**Figure 1:**
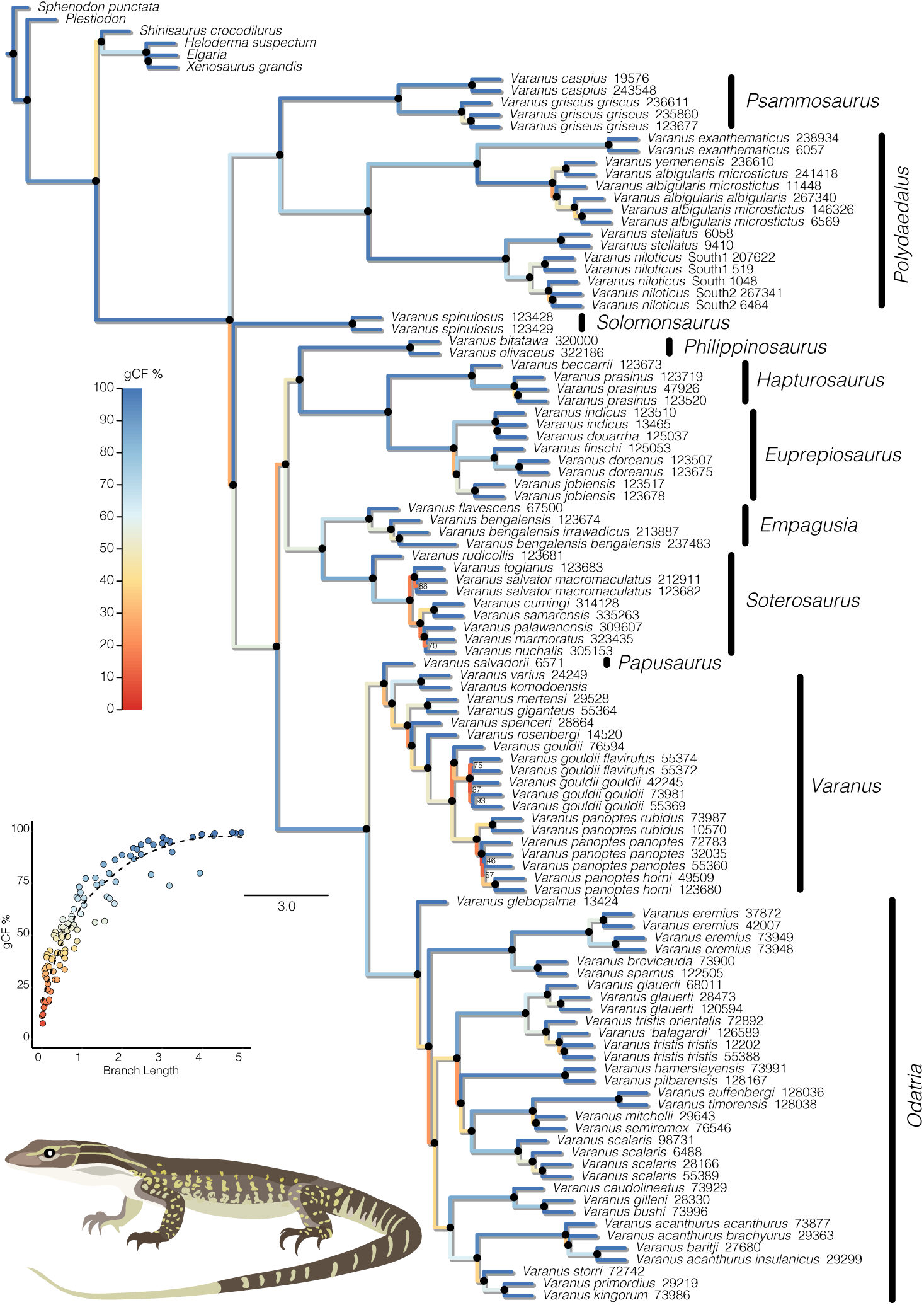
The fully sampled species tree estimated with ASTRAL is largely concordant with our total evidence species tree (Fig.S3). Nodes denoted by a *•* black circle are supported by local posterior probability values >0.90, all others (<0.90) are considered equivocal and designated by lpp values. Branch colors correspond to gene concordance factors, and represent the percent of gene trees which decisively support the presented bifurcation. Inset plot shows that as expected, gCF values increase with increasing branch lengths, shown in coalescent units. Subgeneric names are listed to the right of each group.

Overall, we suggest a younger timeline for the diversification of modern varanid lizards when compared to other phylogenetic studies, with a crown age in the early-to-mid Oligocene. This timing suggests *Varanus* potentially dispersed into the Indo-Australian region shortly after the collision of the Australian and Asian plates. If this is true, the connection of Sahul to Sundaland likely facilitated the dispersal of monitor lizards across an Indonesian island bridge, and extensive over-water dispersals seem less probable. Similarly, this proximity has also allowed small Australopapuan *Varanus* like the *V. scalaris* complex, as well as the largest extant monitor *V. komodoensis* to disperse back into the Indonesian archipelago (at least Wallacea). This pattern is consistent with the adaptive radiation of Australopapuan elapid snakes (Keogh 1998) and pythons (Reynolds et al. 2014; Esquerre et al. 2019), from Asian origins, and may underlie a more common diversification trend.

### Competition, Character Displacement & Size Evolution

Despite a relatively conservative body form, *Varanus* lizards have diverged into a number of ecologies and an astonishing array of body sizes. These include highly crytpozoic pygmy monitors like *V. primordius*, slender canopy dwellers like *V. prasinus*, the stout-bodied semiaquatic *V. mertensi* and *V. salvator* complex, and monstrous apex predators like the Komodo dragon *V. komodoensis*and extinct *V. priscus*. Across their range, monitors have also converged ecomorphologically with a number of mammalian predators, potentially putting them in direct competition for resources (Sweet and Pianka 2007). Competition is expected to influence interacting lineages by driving similar organisms apart in geographic space (exclusion), or in phenotypic or behavioral traits (character displacement) (Brown and Wilson 1956). In Australia, the diversity of varanids is matched by that of carnivorous marsupials, which vary from tenacious shrew-sized ningauis (*Ningaui*) up to the recently extinct wolf-like *Thylacine*.

By modelling the evolution of body size of Australian monitors and dasyuromorph and peramelemorph marsupials using lineage interaction-informed PCMs, we find strong support for the accumulation of size disparity as a result of character displacement independently and in parallel in these two groups. This is corroborated by greater than expected functional diversity of monitor assemblages (over dispersion). However, we do not find evidence of competition between marsupials and monitors and instead size evolution appears to have been dictated instead by within-clade character displacement. This may seem counterintuitive, considering carnivorous marsupials and monitors largely overlap in diet and size, with small animals—monitors and marsupials alike—eating large invertebrates and small lizards, and larger animals taking larger vertebrate prey (James et al. 1992). But, marsupial predators and monitors differ in one very basic way, which is their activity period. Both are active foragers, covering wide tracks of land in search of food, but while monitors are almost exclusively diurnal, often roaming during the hottest part of the day, nearly all faunivorous marsupials are nocturnal. This temporal separation may explain the lack of competition in our analyses, and their continued coexistence. Data from other continents lend some support to this hypothesis. Across Africa, the Indian subcontinent, and throughout Southeast Asia, monitor lizards compete with other diurnal carnivorans, such as herpestids (mongooses), viverrids (civets), canids (dogs), mustelids (weasels), and felids (cats). Throughout these regions, *Varanus* have not diversified to the same extent as in Australia. The possibility of competitive release upon reaching the Australian continent provides a plausible explanation for the diversification of dwarf monitor species (Sweet and Pianka 2007).

While monitor lizards and marsupial predators appear to have diversified without outwardly influencing each others’ trait evolution, both groups appear to have diverged according to character displacement occurring *within* their respective radiations. This suggests that community assembly processes may result in the same observable macroevolutionary patterns across different sympatric groups. Character displacement has long been associated with trait divergence, and was principally described on shallow scales from observable interactions among extant lineages (Vaurie 1951; Brown and Wilson 1956). The practice of extrapolating this idea to fit evolution on geological timescales fits the concept of a micro-to-macro evolutionary spectrum that is dictated by the same processes. The concept of competition as an impetus for evolution however, has been difficult to show explicitly from the fossil or phylogenetic record, and has been criticized for an unnecessarily “progressive” view of the process of evolution (Benton 1987). With the recent development of more appropriate process-generating models, we are now capable of better testing the influence of lineage interactions on evolutionary outcomes (Drury et al. 2016, 2018b; Manceau et al. 2017; Quintero and Landis 2019). In the case of monitor lizards, the exaggerated disparity in body sizes of Australian species is best described by an evolutionary model which accounts for competition among taxa in both space and time. This finding is further supported by evidence of overdispersion in body size variation within monitor communities, suggesting niche partitioning by body size is prevalent across the continent.

## Conclusion

Monitors are an exceptional radiation of lizards capable of traversing sandy deserts and open ocean, living in the canopy and below ground. Here we present a comprehensive phylogenomic hypothesis of the genus, and place them among related varaniform and anguimorph lizards. In agreement with previous study, we find that varanids likely originated in Eurasia in the late Cretaceous or early Paleocene, but have long been spread across Europe, North America, and Africa, with their greatest richness in Indo-Australia. We also present a set of interaction-informed geographically explicit comparative models that help us propose an explanation for the extreme size disparity of living *Varanus*. We suggest that the diversity of sizes of Australian monitors may be the result of a combination of competitive release from carnivorans, and character displacement among other monitor species. Because organisms evolve in natural communities—and not in ecology-free vaccuums—we stress the importance of incorporating macroecological processes into macroevolutionary models. Our methodology involves a stepwise process of estimating ancestral ranges in continuous space (Quintero et al. 2015), then using this to inform interaction matrices in comparative models of trait evolution (Drury et al. 2018b). This framework also provides the opportunity to test the influence of taxa from more than one phylogeny on the evolution of a trait of interest (Manceau et al. 2017), with the goal of better understanding how communities develop and evolve. While our stepwise framework is limited by the unidirectionality of influence (species distributions may dictate trait evolution, but not vice versa), already methods are being developed to jointly infer these processes (Quintero and Landis 2019), as the evolutionary community works to provide a more holistic view of speciation, biogeography, and trait evolution.

## Supplementary Material

### Supplementary Methods

#### Phylogenetic Analyses

To generate a molecular species tree, we started by reconstructing individual genealogies for each of the 388 recovered loci under maximum-likelihood in IQ-TREE (Schmidt et al. 2014). We allowed the program to automatically pick the best fitting model of molecular evolution using PartitionFinder (Lanfear et al. 2012), then perform 1,000 ultrafast bootstraps (Haeseler et al. 2013). As a preliminary step, we also used IQ-TREE to infer the phylogeny from a concatenated alignment, with individual partitions assigned by PartitionFinder. To estimate a species tree, coalescent methods have been shown more accurate than concatenation (Kubatko and Degnan 2007), and so we used the shortcut coalescent method ASTRAL III (Zhang et al. 2017), with all our IQ-TREE gene trees as input. We estimated local posterior probabilities in ASTRAL and gene concordance factors (gCF) to address node support.

As a complementary strategy to estimating *Varanus* relationships using ASTRAL, we also estimated a species tree using the full multispecies coalescent (MSC) model implemented in Computational requirements limit the number of loci we can realistically use under the MSC, and so we summarized per-locus informativeness using AMAS (Borowiec 2016). We then used custom scripts to sort the loci sequentially by (*i*) missing taxa per alignment, (*ii*) number of variable sites, and (*iii*) AT content. Given this order, we then chose the first three sets of twenty loci (1–20; 21–40; 41–60) as representatives of the most informative and complete loci, and used them to build our phylogeny (Fig.S8).

Advances in phylogenetic reconstruction methods have sought to better integrate molecular sequence data with fossil ages and morphological data (Lee et al. 2009; Pyron 2011; Ronquist et al. 2012; Beck and Lee 2014; Heath et al. 2014; Gavryushkina et al. 2017). Incorporating these lines of information in a **combined evidence** approach has provided more accurate phylogenetic estimation, and timing of divergence events. We reconstructed the phylogeny of living and extinct varaniforme lizards using the **Fossilized Birth-Death Multi-Species Coalescent** implemented in starBEAST2 (Ogilvie et al. 2018). In divergence dating analyses fossil information may be included using node priors (generally hard minimum bounds with diffuse upper bounds) or as tip dates (an estimate of the fossil sampling time) (Ho and Phillips 2009). Where data is available, combining node– and tip-dating may provide an advantage over using either method independently (Beck and Lee 2014; O’Reilly and Donoghue 2016). This provides the opportunity to co-estimate the phylogeny and divergence times, while providing structured priors on nodes which may otherwise be driven to unrealistic deep or shallow values. In most implementations of tip-dating fossil ages are fixed to a single value—most often this is the median value between upper and lower bounds. To avoid unintentional bias in choosing exact fossil ages, we instead incorporate uncertainty by sampling from informed uniform priors allowing the fossil ages to be jointly estimated (Barido-Sottani et al. 2019). Morphological data were modelled under the Mkv model, a special case of the Mk model (Lewis 2001)—the most commonly used model for discrete morphological data. The Mk model operates under the assumption that each character may exhibit *k* states, and can transition among states at equal frequencies/rates. Because different characters may exhibit differing numbers of states, we applied the partitioning strategy of Gavryushkina et al. (2017), which partitions the morphological data based on the number of observed states of each character. Traditionally, invariant characters are either not coded, or stripped from discrete morphological alignments, resulting in an ascertainment bias for variable characters. The Mkv model (Lewis 2001) was proposed to account for this. All analyses were run for four independent chains under uncorrelated relaxed lognormal (UCLN) and strict molecular clocks for 1 billion generations and sampled each 5×10^5^ generations, to assess convergence among runs. We inspected the MCMC chains for stationarity (ESS > 200) using Tracer v1.7.0 (Rambaut et al. 2018), and discarded the first 10-40% of each run as burn-in as necessary before combining runs.

Morphological and molecular phylogenies of living and extinct monitor lizards have previously provided conflicting results regarding the relationships between the major clades and subgenera of *Varanus*. Inconsistencies among these data types may partially be due to difficulties in accurately modelling morphological evolution (Goloboff et al. 2018). While our knowledge of the homology, rate, and process of molecular evolution is considerable, it has been much more difficult to adequately model morphological data. In contrast to molecular sites or loci, morphological characters are likely more often correlated (Billet and Bardin 2018), nonhomologous (Baum and Donoghue 2002), or evolving under dramatically different mechanisms (Goloboff et al. 2018), and may disrupt our best efforts at reconstructing phylogeny, divergence times, and rates of evolution. This difficulty is exaggerated on deep time scales and highlights important caveats to consider in the application of combined– or total-evidence methods (Puttick et al. 2017; Luo et al. 2018). To address this, we also estimated divergence dates using an “extant-only” approach, limiting the sampling to living taxa with molecular data, and used the multispecies coalescent model implemented in StarBEAST2. We again used subsets of 20 loci, and applied several node calibrations described in Table S8, and discussed in the Supplemental Material (“Node Priors and *Varanus* in the Fossil Record”). We ran four independent chains under uncorrelated relaxed lognormal (UCLN) clocks with the GTR substitution model applied to all partitions for 1 billion generations and sampled each 5×10^5^ generations, to assess convergence among runs. Again, we inspected the MCMC chains for stationarity (ESS > 200) using Tracer v1.7.0 (Rambaut et al. 2018), and discarded the first 10-20% of each run as burn-in as necessary before combining runs.

#### Fossil Taxa as Sampled Ancestors

Fossil taxa are almost always assumed to represent terminal tips that have since gone extinct. To test this assumption, we allowed fossil taxa to be identified as terminal or stem lineages using the **Sampled Ancestors** package implemented in StarBEAST2. After running our full analyses, we also ran prior-only analyses for each dataset and used these to calculate Bayes factors (BF) for each fossil taxon to test competing hypotheses. Given that we place a prior on the age of each taxon (*τ*) and are jointly estimating their position among the phylogeny, including a model (*M*) of the molecular and morphological evolution, we can sample exclusively from both the prior and posterior of our starBEAST2 analyses (*Supplementary Material*). We used a threshold of log(BF) > 1 to identify sampled ancestors, log(BF) < −1 to recognize terminal taxa, and −1 < log(BF) < 1 taxa were categorized as equivocal.

#### Biogeographic History

*Varanus* lizards have been variously hypothesized to have originated in Asia (Keast 1971; Estes 1983; Fuller et al. 1998; Jennings and Pianka 2004; Amer and Kumazawa 2008; Vidal et al. 2012; Conrad et al. 2012), Africa (Holmes et al. 2010), or Gondwana (Schulte et al. 2003) with conclusions largely based on which taxa were included, and the timing of varanid divergence events. To infer the biogeographic history of varanids and their allies, we used *BioGeoBEARS* (Matzke 2014). Because of the broad distribution of living and extinct monitors, we divided their range into seven major regions relevant to this group: North America, Europe, Sundaland/Wallacea, Australo-Papua, Africa/Arabia, West Asia (Indian subcontinent and surrounds), and East Asia (China, Mongolia, mainland Southeast Asia). We used as input our maximum clade credibility tree from the total evidence dating analysis in order to incorporate the geographic history of fossil taxa. Because of the deep evolutionary history of this group we took plate tectonic history into account by correcting dispersal probability as a function of distance between areas. We estimated distances between areas and continents through time at five million year intervals from 0–40 million years ago, then ten million year intervals from 40–100 million years, using latitude and longitude positions from GPlates (Boyden et al. 2011), and calculated pairwise distance matrices using the R package *geosphere*(Hijmans 2016). Additionally, we limited the model-space by providing information about area adjacency. For each time period, we removed unrealistic combinations of ranges (e.g. North America + AustraloPapua), with the aim of recovering more realistic biogeographic scenarios. We undertake the exercise of reconstructing the biogeographic history of this group fully recognizing that the observation of current (or fossilized) ranges of terminal taxa provide little information about the processes that got them there (Ree and Sanmartín 2018). Recognizing this, we implement only the dispersal-extinction-cladogenesis model (DEC) and the jump extension of this model (DEC+j), and compare models with and without dispersal-distance-penalties. Further, we acknowledge the DEC model’s proclivities for inflating the importance of cladogenetic dispersal, and consider its conclusions cautiously.

To further understand the spatial evolution of *Varanus*, we used a Bayesian method to model the dispersal of monitors across the Australian landscape. The R package *rase* (Quintero et al. 2015) assumes a Brownian motion diffusion process, using point data instead of discrete areas to infer geographic ranges which may be irregular or discontinuous. We started by downloading occurrence records for all continental Australian *Varanus* species from the Atlas of Living Australia (ala.org), curating the data for erroneous records, then trimmed our input tree down to just Australian taxa. We ran *rase* for 10,000 generations, sampling each 10th generation, then discarded the first 10% (100 samples) as burn-in, leaving 900 samples. We inspected the traces of the MCMC chains for stationarity using *coda* (Plummer et al. 2006).

#### Signature of Character Displacement

Ecological communities are generally thought to assemble under opposing processes of habitat filtering and interlineage competition. Filtering is suggested to select for species with similar phenotypes, resulting in conservatism or convergence, whereas competition is expected to result in greater phenotypic disparity. These expectations can be tested by investigating the functional diversity of communities across the landscape. We divided the Australian continent into half-degree cells, and created a site by species matrix using the ALA distribution data for (**i**) monitor lizards and again for (**ii**) monitors and dasyuromorphian marsupials together. We estimated the functional diversity for the two data sets using the package *FD* (Laliberté et al. 2014) and Rao’s Quadratic, using body size as the trait of interest. We then estimated functional diversity for each inhabited cell 100 times using a dispersal null metric model which sampled from nearby cells assuming a probability proportional to the inverse of the distance from the focal cell. To compare observed and simulated functional diversities, we calculated standardized effect sizes (SES) for each cell, and a mean SES across the continent with 95% confidence intervals.

#### Modelling Body Size Evolution with Competition

Only within the past few years have phylogenetic comparative methods (PCMs) begun to account for the interaction of lineages on trait evolution. Building off conceptual work by Nuismer & Harmon (Nuismer and Harmon 2015), Drury et al. (Drury et al. 2016) and Manceau et al. (Manceau et al. 2017) integrated a system of ordinary differential equations in RPANDA (Morlon et al. 2016) for estimating the effect of competition on trait evolution in a maximum likelihood framework. This methodology allows us to estimate a parameter *S* which describes the strength of the interaction, as well as the polarity: negative values of *S* indicate repulsion, positive values indicate attraction towards common values. In its most simplistic form (the Phenotypic Matching **PM**, or Matching Competition **MC** model), the *S* parameter interacts with the mean trait values of all other lineages (vector *X* _t_), to reflect their relationship (*Supplementary Material* Equation 1). To take into account changes through evolutionary time, the *S* parameter further interacts with the evolutionary rate (*σ*), and drift (*d*), to dictate the trajectory of trait evolution. This model however, assumes that *all* lineages in a tree are sympatric and interact with one another. To address this, Drury et al. (Drury et al. 2018b) extended the model by incorporating interaction matrices (*P*) that dictate which taxa interact with one another to more realistically estimate *S* (equation 1).

In natural ecosystems, many different organisms compete for the same resources, so accounting for competition only within a single group is perhaps unrealistic. To address this issue, we consider the influence of another broadly distributed group of like-sized carnivores and omnivores, dasyuromorphian and peramelemorphian marsupials, on the size evolution of Australian monitor lizards. To test this hypothesis we begin by trimming the marsupial phylogeny of Brennan & Keogh (Brennan and Keogh 2018) down to just the faunivorous clade, from which we also dropped *Myrmecobius* because of its unusual ecology. We collected body size (mm) information for marsupials from Pantheria (Jones et al. 2009), and monitors from the literature (Wilson and Swan 2013). Manceau et al. (Manceau et al. 2017) introduced a framework for estimating the effect of one clade on the trait evolution of another, incorporating two phylogenetic trees, referred to as the Generalist Matching Mutualism **GMM** model. This is essentially a two-clade extension of the **PM** model, which makes the assumption that the evolution of trait values in clade A are the result of interactions *only* with lineages in clade B, and vice versa. We present a graphical description of this and additional models below (Fig.S1). The **GMM** model however makes two very basic assumptions that we expect do not fit our data: (**1**) interactions between phenotypes are limited to interclade (between trees) matching or competition, meaning there is no influence of intraclade (within tree) interactions, and (**2**) that all contemporaneous lineages are interacting, regardless of geographic distribution. To address these assumptions, we develop a series of models that expand on the interaction parameter *S*, and incorporate biogeography with the hopes of providing more realistic models of trait evolution. We briefly summarize and illustrate those models here, but discuss their behavior more extensively in the *Supplementary Material*.

Existing and new models described here allow us to test a number of hypotheses regarding the evolution of varanid body size. We focus on those that incorporate dasyuromorphian marsupials as well, because this provides a more holistic view of the macroevolution of two iconic groups of Australian vertebrates. Using these models we first test the idea that the evolution of varanid and dasyuromorphian body size has been dictated by competition with congeners, between clades, or both. We then test whether the strength of intraclade competition is equivalent in the two groups, and if the inclusion of geography via coexistence matrices improves model fit. Finally, we can ask if size evolution is instead dictated by non-ecological processes, by implementing standard models of trait evolution, Brownian Motion **BM** and Ornstein Uhlenbeck **OU**. Using these traditional null models, we can again ask if monitor and dasyuromorphian size has evolved under similar or independent rates using *ratebytree* in *phytools*, though we also provide implementations of shared BM and OU models in the *RPANDA* framework—**CoBM** and **CoOU**.

To incorporate historical and contemporary biogeography, we started by extending our *rase*analyses to marsupials with data collected from the ALA. We designed a number of custom scripts and functions to process the spatial data and model objects including extensions of the ‘CreateGeoObject’ 20 of *RPANDA*. Our functions ‘CreateGeoObject_SP’ and ‘CreateCoEvoGeoObject_SP’ produce *RPANDA* GeoObjects that take as input a tree, spatial distribution data in latitude/longitude format, and a post-processed *rase* object. Internally, these functions use the packages *sp* and *rgeos* to translate spatial data into spatial polygons representative of species distributions. Then, at each cladogenetic event, we determine the pairwise overlap of all contemporaneous lineages to construct our GeoObject (see Fig.S6). The ‘CreateCoEvoGeoObject_SP’ function has adapted this process for two trees, to be applied to GMM-type models.

#### Interaction Model Summaries

The following comparative models are summarized visually in Fig.S1.

The *Phenotypic Matching* (*PM*) (Nuismer and Harmon 2015) or *Matching Competition* (*MC*) (Drury et al. 2016) model is the basis for many PCMs incorporating interactions between lineages. *S* is estimated from the interaction of *all* contemporaneous lineages.

The *Phenotypic Matching Geography* (*PM_geo_* or *MC_geo_*) (Drury et al. 2016) model was built as a geographic extension to the PM/MC model. Originally designed to account for sympatric island lineage interactions (determined in *P* matrices), only codistributed species influence the estimation of *S*.

The *Generalist Matching Mutualism* (*GMM*) model. Assumes equal interaction (S) between all inter-clade lineages, but no interaction (0) among lineages within a tree (intra-clade). We embrace a broad description of the GMM mdoel, where *S* can be positive indicating attraction towards the mean trait value of interacting lineages, or negative indicating repulsion away from the mean trait value of interacting lineages. Because interactions are estimated only **between** clades, *p_k,l_*=1 if lineages *k* and *l* are from different clades (trees), and *p_k,l_*=0 for any two lineages *k* and *l* from the same clade (tree).

The *Generalist Matching Mutualism All* (*GMM_all_*) model. Assumes equal interaction (S) between all taxa in both trees (inter- and intra-clade). *S* can be positive indicating attraction towards the mean trait value of interacting lineages, or negative indicating repulsion away from the mean trait value of interacting lineages, but *p_k,l_*=1 always.

The *CoEvo* model. An extension of the GMM model, accounting for interactions only between geographic co-occurring lineages. As with the GMM model, it only estimates interaction (S) between taxa across trees (inter-clade, *not* intra-clade). This model also properly accounts for the number of co-occuring lineages by dividing S (Pk/l) using rowsums (see Manceau et al. pg.559, equation 7).

The *CoEvo_all_* model. This is a CoEvo extension of the GMM_all_ model, estimating interaction (*S*) between all co-occurring taxa (inter-clade *and* intra-clade). *p_k,l_*=1 always. In calculating the interaction matrices, this model accounts for the number of co-occuring lineages by dividing *S* /*p_k,l_*, assuming an equal strength of interaction with each cohabiting lineage.

The *CoEvo_split_* model. Again, an extension of the GMM model, accounting for interactions only between geographic co-occurring lineages. It accounts for interactions between all taxa like the CoEvo_all_ model, but estimates a different interaction parameter for intra-clade (S2) and inter-clade (S1) interactions. In calculating the interaction matrices, this model accounts for the number of co-occuring lineages by dividing *S* /*p_k,l_*, assuming an equal strength of interaction with each cohabiting lineage. This model is identical to the models: CoPM_geo_ if *S* _1_=0, CoEvo if *S* _2_=0, and CoEvo_all_ if *S* _1_=*S* _2_.

The *CoPM* model. This is a joint estimation of the PM model for two trees. It estimates single interaction (*S*) and rate (*σ*) values for both trees, but *S* is estimated solely from intra-clade interactions (no interaction between trees). All lineages in a tree are assumed to interact with *all*other lineages in that tree

The *CoPM_geo_* model. This is an extension of the CoPM model, which is a joint estimation of the PM model for two trees. It estimates single interaction (*S*) and rate (*σ*) values for both trees, but *S* is estimated solely from intra-clade interactions (no interaction between trees). It correctly accounts for interaction only among geographic overlapping lineages, and corrects the interaction estimate for the number of overlapping lineages.

The *JointPM* model. This is a joint estimation of the PM model for two trees. It differs from the CoPM model by estimating separate interaction values for each clade (tree_1_ = *S* _1_; tree_2_ = *S* _2_). All lineages in a tree are assumed to interact with *all* other lineages in that tree.

The *JointPM_geo_* model. This is a joint estimation of the PM model for two trees. It differs from the CoPM_geo_ model by estimating separate interaction values for each clade (tree_1_ = *S* _1_; tree_2_ = *S* _2_). Like the CoPM_geo_ (unlike JointPM) it correctly estimates the interaction parameters (*S* _1_,*S* _2_) for only geographic overlapping taxa (it also corrects for the number of taxa overlapping). This model is identical to the CoPM_geo_ model if *S* _1_=*S* _2_.

#### Model Behavior and Identifiability

The ability to identify competition and estimate associated parameters using process-based models has been tested extensively previously (Drury et al. 2016, 2018a, 2018b). From this we know that the ability to recover competitive models and estimate the interaction parameter *S* —when it is the generating process—is strongly linked to the absolute value of *S*, and to a lesser degree the size of the phylogeny. Parameter estimate and recovery of *S* can also be highly influenced by the incorporation of stabilizing selection (*ψ* or *α*), with the two parameters working agonistically in instances of competition (-*S*), and synergistically in mutualistic circumstances (+*S*).

To ensure that we can accurately identify our models and estimate parameter values, we undertook a focused simulation exercise. Following the advice of Manceau et al. (Manceau et al. 2017), we simulated data directly onto our Australian monitor and marsupial trees under the same models we fit to our empirical data: BM_shared_, OU_shared_, CoEvo, CoEvo_all_, CoEvo_split_, JointPM_geo_, and CoPM_geo_. We used the *RPANDA* function ‘simulateTipData’ to simulate body size data under all specified models, keeping the empirical biogeography constant. Specifics of the generating parameter values are noted in the Table S3. We then iteratively fit the models to our simulated data, and compared fit using AICc and plotted AICc weights. To determine the ability to accurately recover parameter values, we then compared estimated to simulated values under each model.

#### Historical Models of Monitor Size Evolution

To test our hypothesis of character displacement as a driving force of *Varanus* size disparity, we also fit standard stochastic (Brownian Motion) and stabilizing (Ornstein-Uhlenbeck–OU) models of trait evolution, and a multi-optima (OUM) model following Collar et al. (2011). This multi-OU (OUM) model explains size evolution as a result of differing selective optima correlated with habitat use. These models were implemented and fit using *geiger* (Pennell et al. 2014) and *OUwie* (Beaulieu et al. 2012).

Equation 1. Following Manceau et al. (2017), we can estimate if lineage *k* is repelled from (-*S*) or attracted to (+*S*) the average trait value of the lineages it interacts with. *S* represents the the *strength* of the interaction on trait evolution. *d*_1_ and *d*_2_ represent the shift values for lineages from clade 1 and 2 respectively, with the expectation that *d*_1_+*d*_2_=0. *δ_k_* equals one if lineage *k* belongs to clade 1, and zero it if belongs to clade 2, and *p_k,l_* equals one if lineages *k* and *l* interact (in our case it is assumed if they are sympatric) and zero otherwise. *n_k_* = Σ_*l*_*p_k,l_* is the number of lineages interacting with lineage *k*, and *n* is the total number of lineages.

**Table S1.**
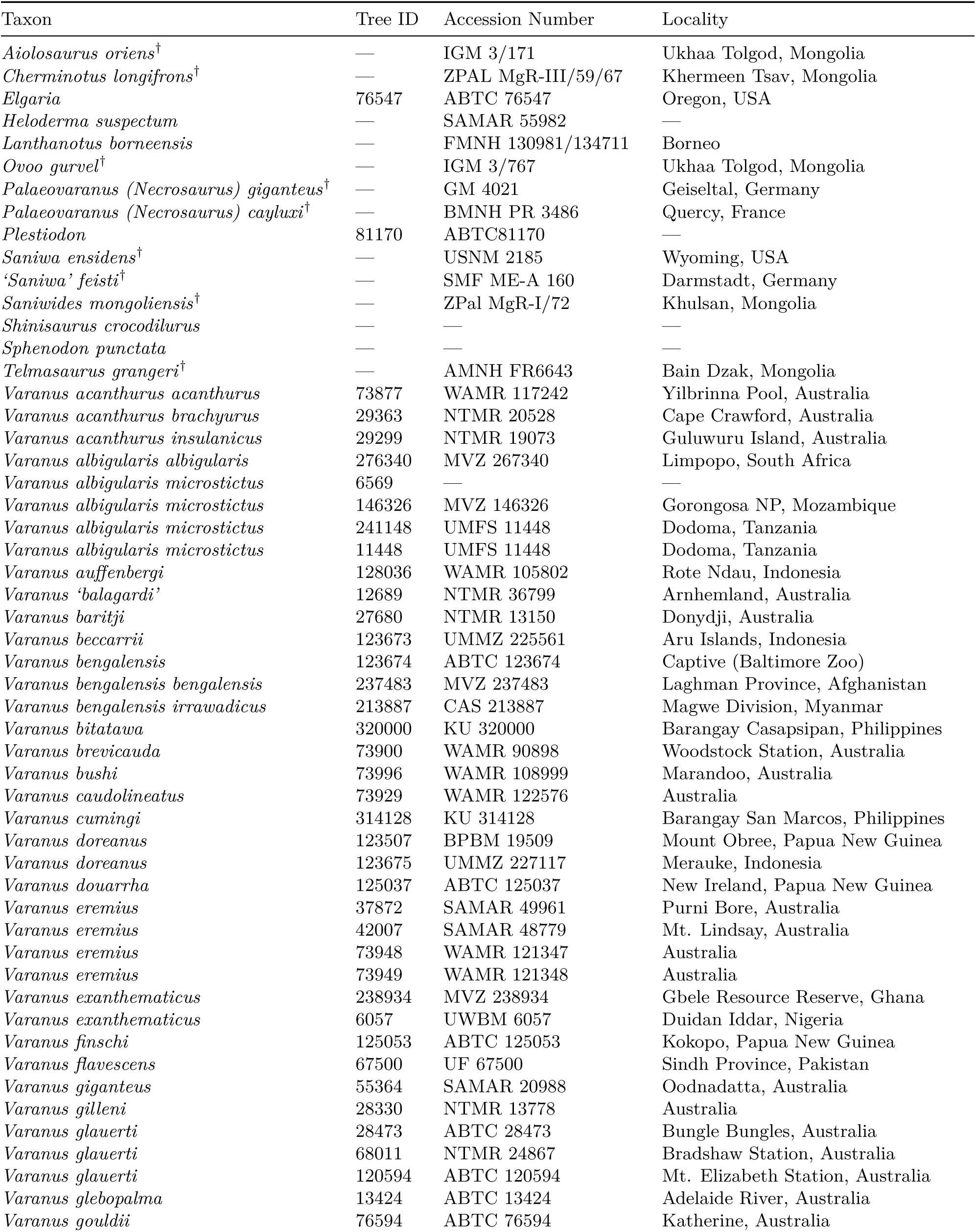
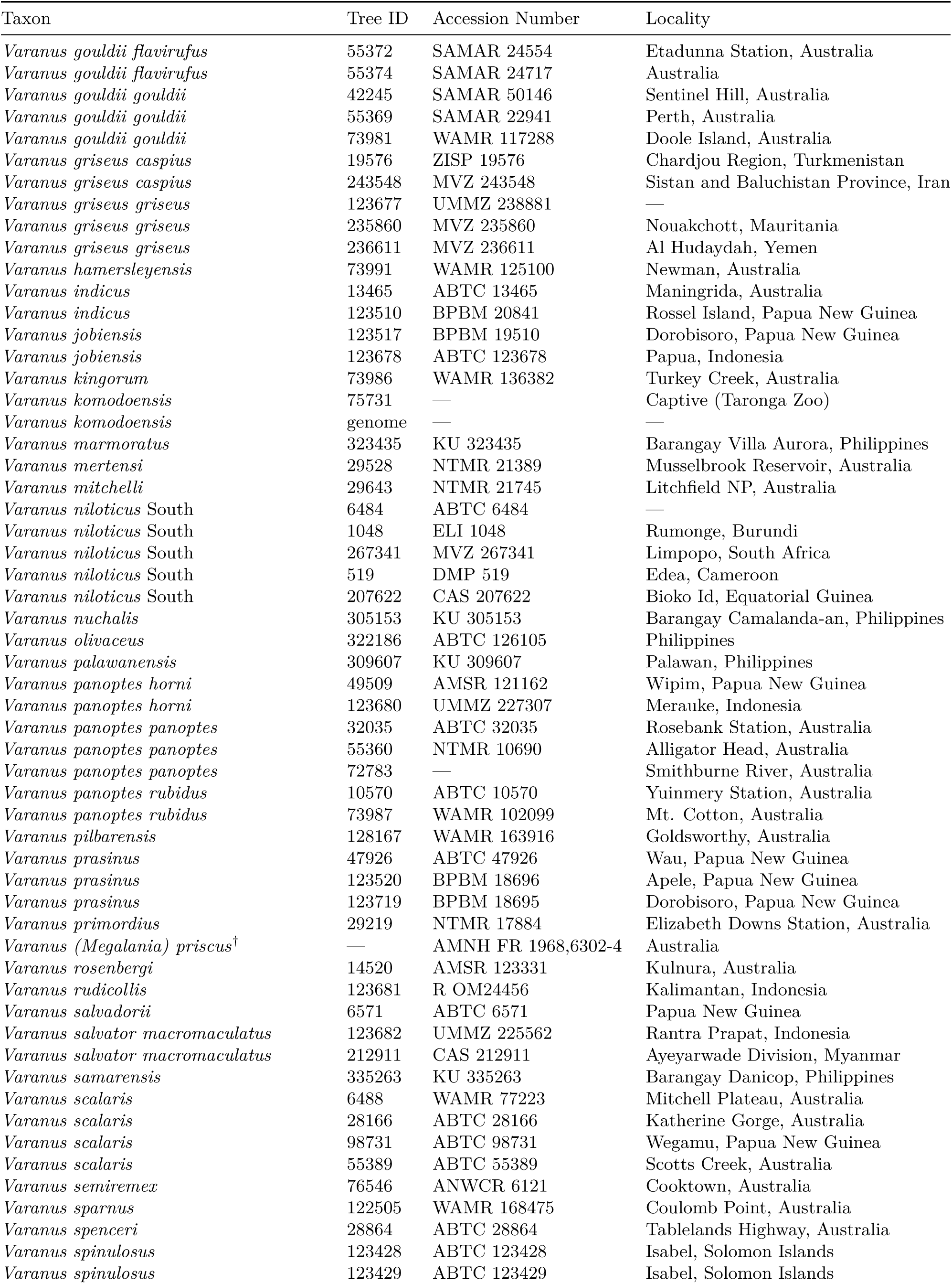
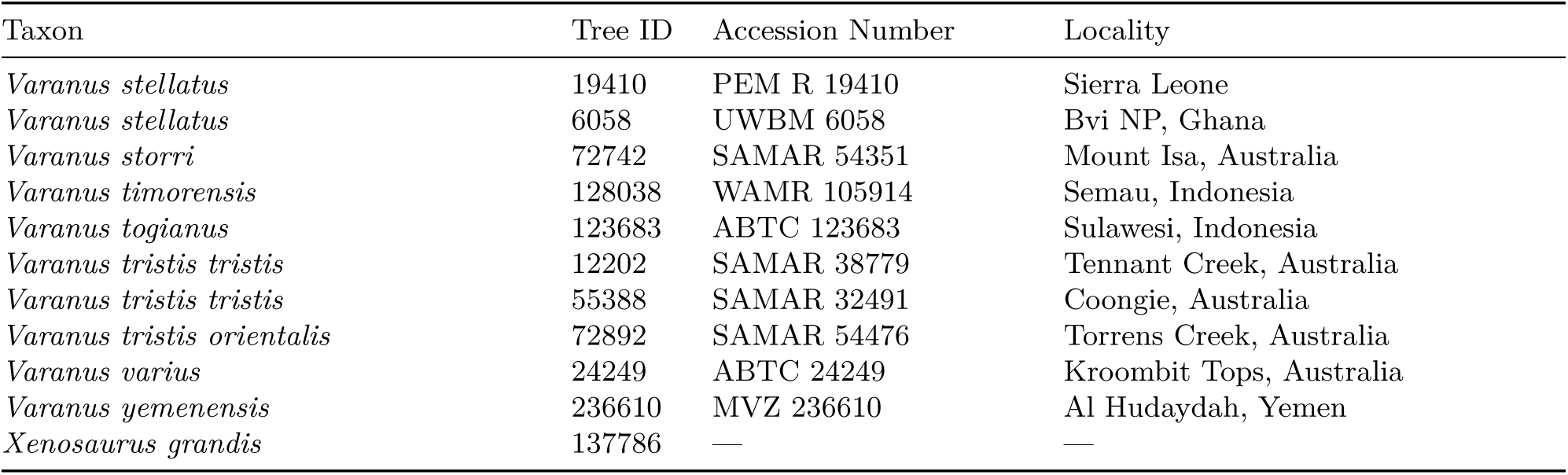
Taxon sampling for this project. The † symbol denotes extinct taxa included in the combined evidence analyses. *Lanthanotus borneensis* is the only extant taxon lacking molecular data.

**Table S2.**
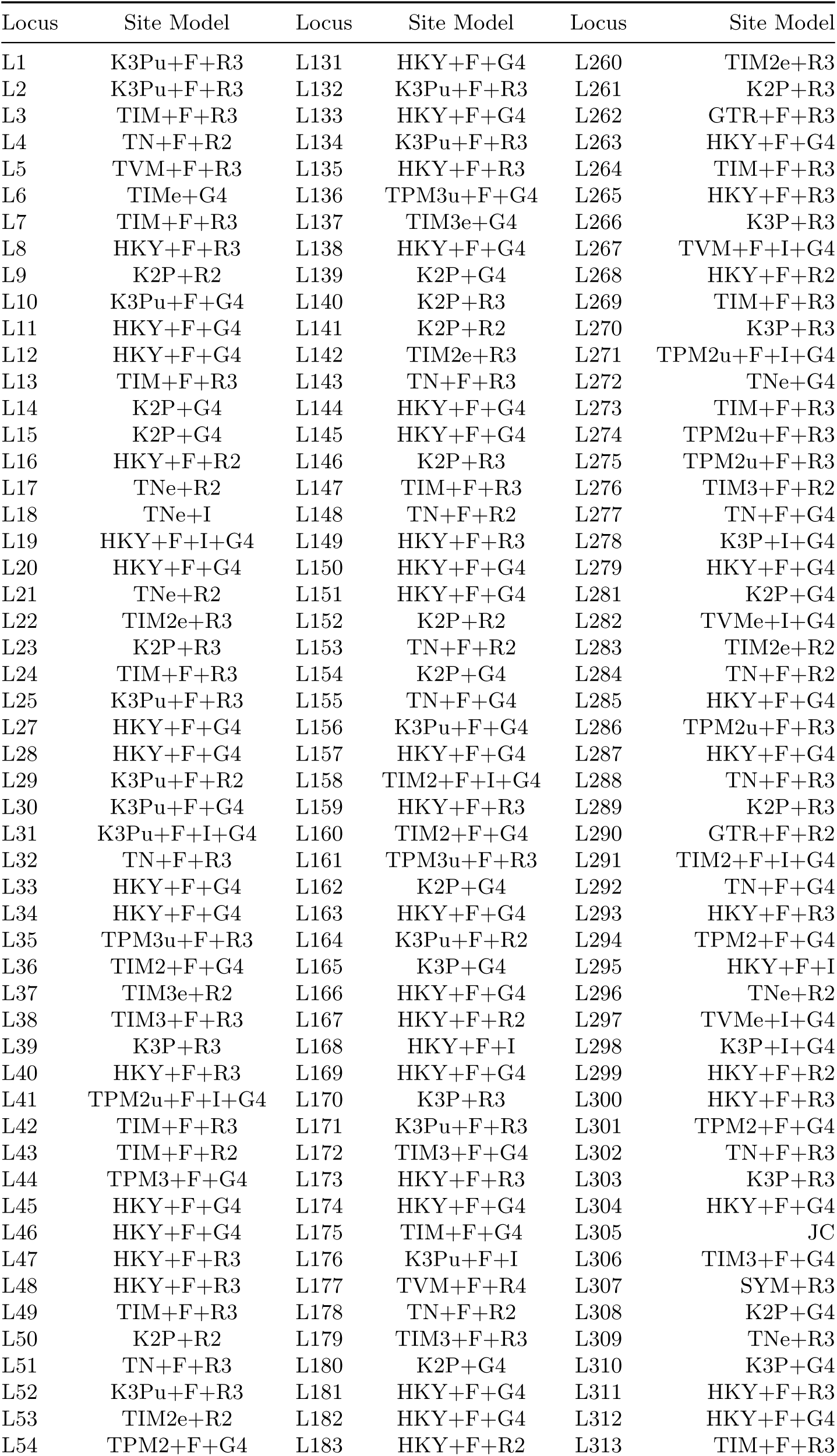
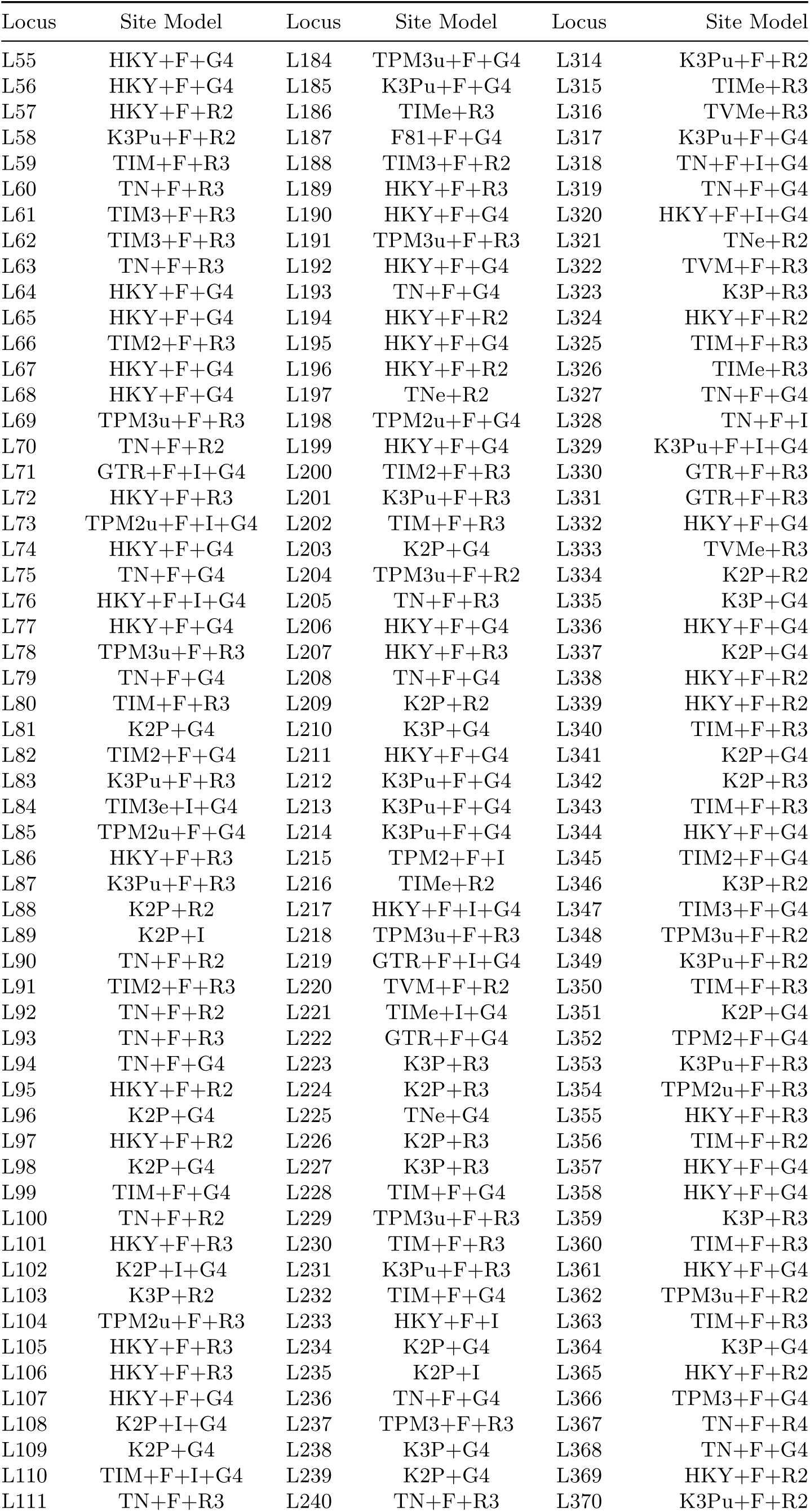
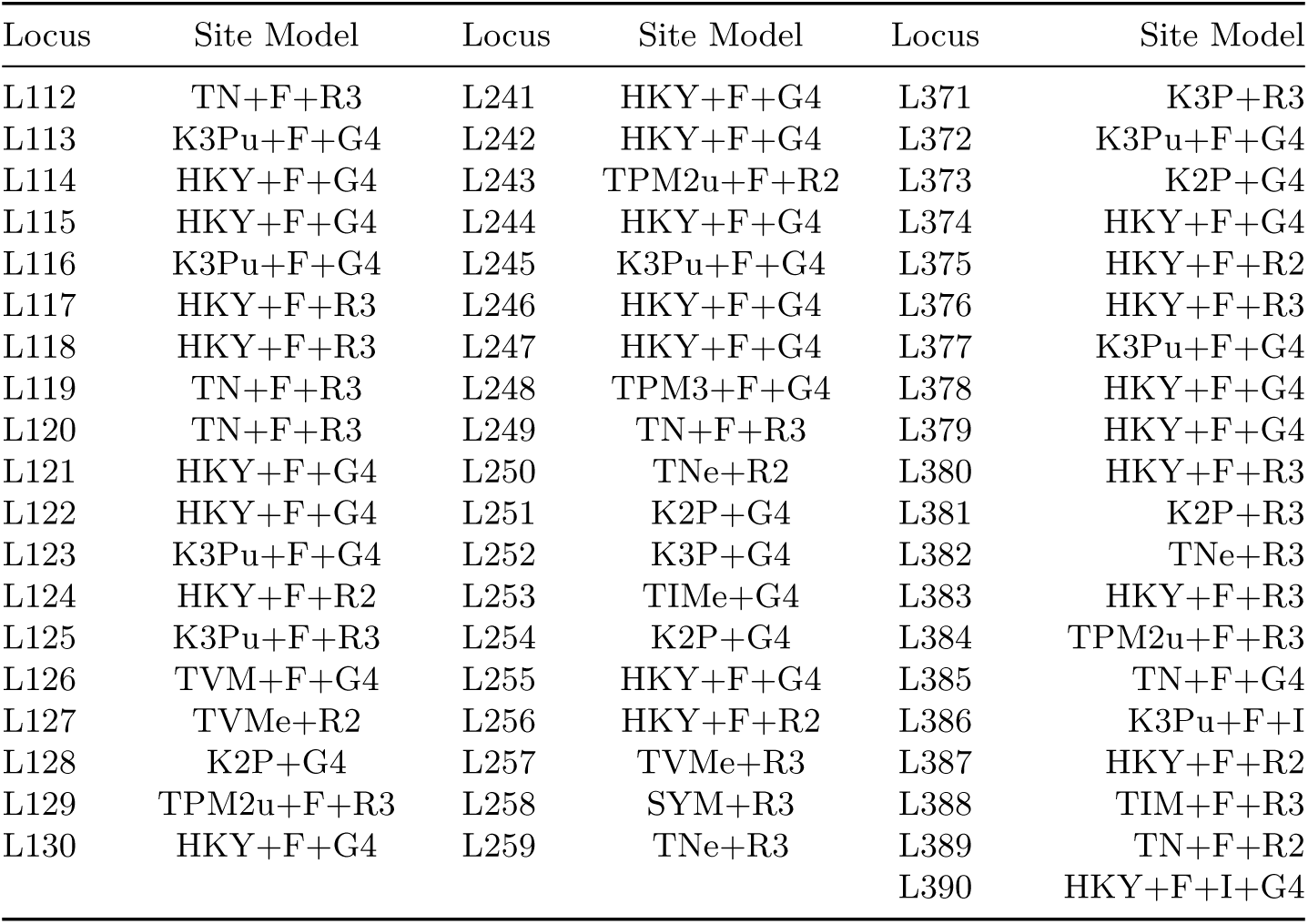
Per locus best fitting models of molecular evolution, determined by IQ-TREE and the Bayesian Information Criterion (BIC). Independent gene trees were estimated using the preferred model, and 1,000 ultrafast bootstraps.

#### Disruptive Morphological Samples

We removed several morphological samples because of disruptive RogueNaRok scores in our preliminary analyses. Those taxa and their respective improvement values are listed below. Additional fossil taxa (*V. mytilini, V. marathonensis, V. hooijeri, V. cf. bengalensis*) were removed from the alignments *before* running combined evidence analyses becasue of their highly fragmentary and unstable phylogenetic nature.

**Table S3.**
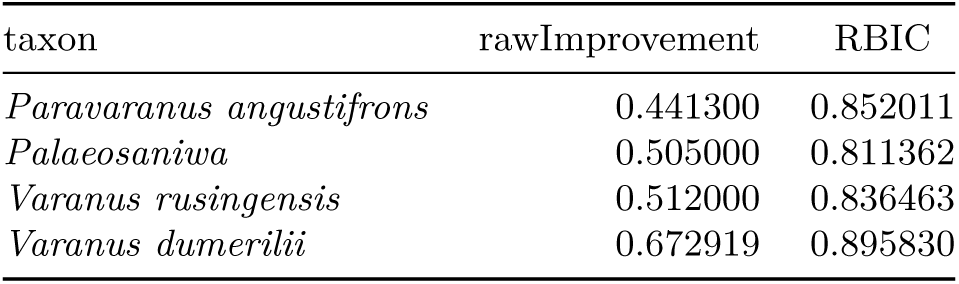
RogueNaRok scores for disruptive samples, which were ultimately pruned from final dating analyses.

#### Simulation Exercise

**Table S4.** We simulated traits onto our empirical trees under the parameters below.

| Model | $\sigma$ | $\alpha$ |
| --- | --- | --- |
| Brownian Motion | 0.003, 0.03, 0.3 | 0 |
| Ornstein-Uhlenbeck | 0.3 | 0.03, 0.06, 0.12 |

| Model | $m_0$ | $v_0$ | $d_1$ | $d_2$ | $\sigma$ | $S_1$ | $S_2$ |
| --- | --- | --- | --- | --- | --- | --- | --- |
| CoEvo | 0 | 0 | -0.01 | 0.01 | 0.3 | -0.001, -0.01, -0.1, -1 | — |
| CoEvo<sub>all</sub> | 0 | 0 | -0.01 | 0.01 | 0.3 | -0.001, -0.01, -0.1, -1 | — |
| CoPM<sub>geo</sub> | 0 | 0 | -0.01 | 0.01 | 0.3 | -0.001, -0.01, -0.1, -1 | — |
| JointPM<sub>geo</sub> | 0 | 0 | -0.01 | 0.01 | 0.3 | -0.001, -0.01, -0.1, -1 | -0.005, -0.05, -0.5, -5 |
| CoEvo<sub>split</sub> | 0 | 0 | -0.01 | 0.01 | 0.3 | -0.001, -0.01, -0.1, -1 | -0.005, -0.05, -0.5, -5 |

#### Empirical Model Fitting

**Table S5.** Model fitting results to accompany Figure 4.

| Model | logLik | Param. | AIC | AICc | deltaAICc | AICcWt |
| --- | --- | --- | --- | --- | --- | --- |
| CoPM<sub>geo</sub> | -23.90383 | 6 | 59.80767 | 60.78441 | 0.000000 | 0.484718461 |
| JointPM<sub>geo</sub> | -23.73944 | 7 | 61.47888 | 62.79653 | 2.012117 | 0.177240890 |
| CoEvo<sub>Split</sub> | -23.83223 | 7 | 61.66447 | 62.98212 | 2.197701 | 0.161534338 |
| CoEvo<sub>all</sub> | -25.49440 | 6 | 62.98880 | 63.96554 | 3.181127 | 0.098790805 |
| BM<sub>shared</sub> | -29.90901 | 3 | 65.81802 | 66.08768 | 5.303269 | 0.034190017 |
| CoEvo | -27.22316 | 6 | 66.44632 | 67.42307 | 6.638654 | 0.017535744 |
| BM<sub>ind</sub> | -29.77132 | 4 | 67.54264 | 67.99719 | 7.212771 | 0.013160008 |
| OU<sub>shared</sub> | -29.90844 | 4 | 67.81688 | 68.27142 | 7.487007 | 0.011473783 |
| OU<sub>ind</sub> | -29.78290 | 6 | 71.56580 | 72.54254 | 11.75812 | 0.001355954 |

**Table S6.** Model fitting results to accompany Figure S12.

| Model | logLik | Param. | AIC | AICc | deltaAICc | AICcWt |
| --- | --- | --- | --- | --- | --- | --- |
| CoPM<sub>geo</sub> | -23.90383 | 6 | 59.80767 | 60.78441 | 0.0000000 | 0.2750436948 |
| CoPM | -24.00661 | 6 | 60.01322 | 60.98997 | 0.2055527 | 0.2481798347 |
| JointPM<sub>geo</sub> | -23.73944 | 7 | 61.47888 | 62.79653 | 2.0121169 | 0.1005717613 |
| JointPM | -23.81588 | 7 | 61.63177 | 62.94942 | 2.1650026 | 0.0931702707 |
| CoEvo<sub>Split</sub> | -23.83223 | 7 | 61.66447 | 62.98212 | 2.1977010 | 0.0916593953 |
| GMM<sub>all</sub> | -25.28098 | 6 | 62.56195 | 63.53870 | 2.7542820 | 0.0693932078 |
| CoEvo<sub>all</sub> | -25.49440 | 6 | 62.98880 | 63.96554 | 3.1811274 | 0.0560568457 |
| GMM | -26.43762 | 6 | 64.87524 | 65.85199 | 5.0675740 | 0.0218268965 |
| BM<sub>shared</sub> | -29.90901 | 3 | 65.81802 | 66.08768 | 5.3032690 | 0.0194004344 |
| CoEvo | -27.22316 | 6 | 66.44632 | 67.42307 | 6.6386539 | 0.0099503035 |
| BM<sub>ind</sub> | -29.77132 | 4 | 67.54264 | 67.99719 | 7.2127713 | 0.0074673808 |
| OU<sub>shared</sub> | -29.90844 | 4 | 67.81688 | 68.27142 | 7.4870071 | 0.0065105660 |
| OU<sub>ind</sub> | -29.78290 | 6 | 71.56580 | 72.54254 | 11.758126 | 0.0007694085 |

**Table S7.** Model fitting results for *Varanus* only analyses. The PMOU_geo_ model is identical to the PM_geo_ model, but without the alpha/psi parameter of the OU process, meaning traits are not constrained to evolve around a single optimum value.

| Model | logLik | Param. | AIC | AICc | deltaAICc | AICcWt |
| --- | --- | --- | --- | --- | --- | --- |
| PMOU<sub>geo</sub> | -6.503810 | 4 | 21.00762 | 22.54608 | 0.000000 | 0.67565891 |
| BM | -7.963686 | 4 | 23.92737 | 25.46583 | 2.919753 | 0.15693186 |
| OU | -7.367979 | 5 | 24.73596 | 27.13596 | 4.589877 | 0.06808452 |
| PM<sub>geo</sub> | -5.955623 | 6 | 23.91125 | 27.41125 | 4.865166 | 0.05932944 |
| OUM | -6.349967 | 6 | 24.69993 | 28.19993 | 5.653854 | 0.03999527 |

**Table S8.** Node and tip prior information. Bolded clades indicate priors which were constrained to be monophyletic. In our taxon sampling: **Lepidosauria** represents the root (all taxa—Rhynchocephalians + Squamata); **Anguimorpha** comprises all taxa to the exclusion of *Sphenodon* and *Plestiodon*; Neoanguimorpha comprises *Heloderma*, *Xenosaurus*, and *Elgaria*; *Saniwa ensidens* represents the split between *Lanthanotus* and *Varanus*; and *Varanus rusingensis* is the oldest discernible crown *Varanus*. An alternative node calibration for the crown of extant monitor lizards, the “Jebel Qatrani” *Varanus* (Holmes et al. 2010), is included here, but used only to illustrate the influence of such a calibration (Fig.S4).

| Fossil Taxon | Min.Age | Max.Age | Clade | Calibration |
| --- | --- | --- | --- | --- |
| Vellberg Jaw | 238 | — | **Lepidosauria** | Exp.; M=10, Off=238 |
| *Dalinghosaurus* | 113 | — | **Anguimorpha** | Log; M=2, S=1, Off=113 |
| *Primaderma* | 98 | — | Neoanguimorph | Exp.; M=4, Off=98 |
| *Saniwa ensidens* | 48 | — | *Lanthanotus* + *Varanus* | Gamma; A=2, B=8, Off=45 |
| *Varanus rusingensis* | 16 | — | Crown ***Varanus*** | Gamma; A=2, B=8, Off=16 |
| *Varanus komodoensis* | 3.6 | — | *V. komodoensis* + *V. varius* | Exp.; M=1, Off=3.6 |
| Jebel Qatrani *Varanus* | 30 | — | Crown ***Varanus*** | Gamma; A=2, B=8, Off=30 |
| ... | ... | ... | ... | ... |
| *Aiolosaurus oriens* | 70.6 | 84.9 | tip | Uniform; min-max |
| *Cherminotus longifrons* | 70.6 | 84.9 | tip | Uniform; min-max |
| *Ovoo gurvel* | 70.6 | 84.9 | tip | Uniform; min-max |
| *Palaeovaranus cayluxi* | 33.9 | 37.2 | tip | Uniform; min-max |
| *Palaeovaranus giganteus* | 33.9 | 48.6 | tip | Uniform; min-max |
| *Saniwa ensidens* | 50.3 | 48.8 | tip | Uniform; min-max |
| *‘Saniwa’ feisti* | 40.4 | 48.6 | tip | Uniform; min-max |
| *Saniwides mongoliensis* | 70.6 | 84.9 | tip | Uniform; min-max |
| *Telmasaurus grangeri* | 70.6 | 84.9 | tip | Uniform; min-max |
| *Varanus priscus* | 0.012 | 0.126 | tip | Uniform; min-max |

#### Node Priors and *Varanus* in the Fossil Record

Monitor lizards and their relatives are not rare in the fossil record, however the phylogenetic affinities of fossil taxa have been difficult to resolve. This is perhaps best captured by Ralph Molnar in his chapter titled *The Long and Honorable History of Monitors and their Kin* (Molnar 2004):

> “Although some of the Cretaceous monitors, particularly those from Mongolia, are known from nice skulls, words like ‘fragmentary’ and ‘frustrating’ involuntarily spring to mind when considering the fossil record of varanids, particularly of *Varanus* itself.”

There is a relatively large resource of fossil *Varanus* material. Many of these fossils have been identified in the literature, however comparatively few have been assigned to living or extinct species, and even fewer have been scored and included in phylogenetic analyses. This makes the inclusion of this fossil information difficult. We quickly discuss how some known and other rumored fossils could potentially be used to date the diversification of monitor lizards, but admit this is nowhere near a complete library of fossil varanids and defer to Molnar’s Molnar (2004) publication for a more thorough discussion of fossil *Varanus*.

To calibrate our phylogeny, we used a combination of node and tip priors to incorporate fossil taxa that were directly sampled (tips) in morphological data or indirectly sampled (nodes) using estimated fossil ages. Previous studies of monitor lizards have used varied calibration schemes to estimate divergence times. The most influential of these has been the application of a hard minimum prior on the crown age of *Varanus* (Vidal et al. 2012; Portik and Papenfuss 2012). This minimum bound is either attributed to the age of the ‘Jebel Qatrani *Varanus*’ (Holmes et al. 2010), or the ‘Yale Quarry *Varanus*’ (Smith et al. 2008). However, based on preliminary assessment (Smith et al. 2008), and more extensive morphological analyses of monitors and their kin (Conrad et al. 2012; Ivanov et al. 2017), these fragmentary fossils are not recovered within the crown of *Varanus* and are more likely stem varanids, suggesting that they should not be used to constrain the minimum age of extant *Varanus*.

In Australia, Stirton (1961) mentioned varanid material from the Etadunna formation, however this material was misattributed, and appears to actually have been a snake (Estes 1984). Estes (1984) further went on to briefly discuss the existence of *Varanus* fossil material from the Mid Miocene Lake Ngapakaldi area, though these vertebrae have not since been described. The same goes for Oligo-Miocene material mentioned by Scanlon (2014), which comes from the Hiatus and White Hunter sites of Riversleigh World Heritage Area (Scanlon 2014). Interestingly, the Riversleigh material contains “the occasional isolated tooth, jaw element, or limb or girdle bone” in addition to the more common vertebrae. Hiatus and White Hunter sites have been dated via biocorrelation (15–25 ma), but could not be radiometrically dated (Woodhead et al. 2016). Also mentioned by Scanlon (2014) are Miocene fossils from Bullock Creek and Alcoota, which are roughly 11-16 ma and 5-12 ma respectively (Murray & Megirian, 1992), but have not been described, evaluated, or scored. Many of these fossils would be particularly valuable for dating the Australian radiation of *Varanus*, but again cannot be placed within the crown of Australian *Varanus* (*Odatria* + *Varanus*), and so should probably only be used to provide a minimum age on the divergence between the Australian radiation and the Asian clade (*Soterosaurus*, *Empagusia*, *Euprepriosuarus*, *Hapturosaurus*, et al.).

Other more recent and perhaps applicable fossils were described by Hocknull (2009) and include a number of cranial and postcranial elements from both *Varanus komodoensis* and an extinct taxon from Timor. The oldest material ascribable to *V. komodoensis* are from Early Pliocene sites at Bluff Downs in northeastern Queensland, Australia. These sites have been dated using whole rock K/Ar (potassium-argon) methods of the overlaying basalt layer to 3.6 million years old. Using this informationw we can set a minimum prior on the divergence between *V. komodoensis* and its closest living relative *V. varius* of 3.6 ma. We outline the remaining node calibrations we applied in our analyses in Table S8.

**Table S9.** Standard effect sizes across Australian *Varanus* communities of varying richness.

| CI Lower | CI Upper | Error | Mean | SD | N | Richness |
| --- | --- | --- | --- | --- | --- | --- |
| 0.01765771 | 0.12753556 | 0.05493893 | 0.07259663 | 0.9740691 | 1210 | 2–11 (all) |
| -0.25265258 | -0.08109514 | 0.08577872 | -0.1668738 | 1.0689687 | 599 | 2 |
| 0.15781727 | 0.36964206 | 0.10591239 | 0.26372966 | 0.8588049 | 255 | 3 |
| 0.24372979 | 0.49436858 | 0.12531940 | 0.36904918 | 0.8026238 | 160 | 4 |
| 0.22269865 | 0.48620457 | 0.13175296 | 0.35445161 | 0.5996291 | 82 | 5 |
| 0.19333878 | 0.71750885 | 0.26208504 | 0.45542381 | 0.9877488 | 57 | 6 |
| -0.00146655 | 0.53272406 | 0.26709530 | 0.26562875 | 0.6612764 | 26 | 7 |
| -0.13678194 | 0.37627128 | 0.25652661 | 0.11974467 | 0.4442921 | 14 | 8 |
| -0.28734231 | 0.21086400 | 0.24910315 | -0.0382391 | 0.3482222 | 10 | 9 |
| -0.97490634 | 0.32296756 | 0.64893695 | -0.3259693 | 0.5226349 | 5 | 10 |
| -1.65996651 | 1.56567795 | 1.61282223 | -0.0471442 | 0.1795088 | 2 | 11 |

**Table S10.**

| CI Lower | CI Upper | Error | Mean | SD | N | Richness |
| --- | --- | --- | --- | --- | --- | --- |
| -0.5327245 | -0.44379796 | 0.04446327 | -0.4882612 | 3 0.9785372 | 1863 | 2–17 (all) |
| -0.9799777 | -0.70273700 | 0.13862037 | -0.8413573 | 7 1.3797493 | 383 | 2 |
| -0.6111262 | -0.39385028 | 0.10863795 | -0.5024882 | 2 1.0244188 | 344 | 3 |
| -0.4704297 | -0.27335183 | 0.09853892 | -0.3718907 | 4 0.8643596 | 298 | 4 |
| -0.3990601 | -0.20963823 | 0.09471094 | -0.3043491 | 7 0.7557160 | 247 | 5 |
| -0.5503313 | -0.34245597 | 0.10393767 | -0.4463936 | 4 0.7145822 | 184 | 6 |
| -0.4627334 | -0.23970379 | 0.11151482 | -0.3512186 | 0 0.6324771 | 126 | 7 |
| -0.4751302 | -0.14600575 | 0.16456224 | -0.3105680 | 0 0.7250327 | 77 | 8 |
| -0.4569476 | -0.17121931 | 0.14286415 | -0.3140834 | 6 0.5625626 | 62 | 9 |
| -0.5162656 | -0.14677388 | 0.18474588 | -0.3315197 | 6 0.6568637 | 51 | 10 |
| -0.7607036 | -0.25664101 | 0.25203131 | -0.5086723 | 2 0.7107791 | 33 | 11 |
| -0.9809260 | -0.43973639 | 0.27059482 | -0.7103312 | 0 0.6408206 | 24 | 12 |
| -0.4821964 | 0.07296914 | 0.27758279 | -0.2046136 | 5 0.5398841 | 17 | 13 |
| -1.1456556 | 0.44100276 | 0.79332918 | -0.3523264 | 2 0.7559579 | 6 | 14 |
| -0.9684752 | 0.20328734 | 0.58588127 | -0.3825939 | 2 0.6334908 | 7 | 15 |
| NA | NA | NA | 0.45452696 | NA | 1 | 16 |
| -1.7472785 | 1.61220378 | 1.67974112 | -0.0675373 | 5 0.6761868 | 3 | 17 |

#### Nested Models

We can show that some of the proposed models are nested. We start by simulating data under the simplest intraclade interaction model CoPM_geo_.

~~~
simCoPM_geo <- simulateTipData(modelCoPM_geo, method=1,
                        c(m0=0, v0=0, d1=-0.001, d2=0.001, S=-0.1, sigma=0.1))
~~~

We can then fit the JointPM_geo_, CoPM_geo_, and CoEvo_split_ models, using the *getDataLikelihood* function, keeping all other parameters the same.

~~~
getDataLikelihood(modelCoPM_geo, simCoPM_geo,
                        c(m0=0, v0=0, d1=-0.001, d2=0.001, S=-0.1, sigma=0.1))
getDataLikelihood(modelJointPM_geo, simCoPM_geo,
                        c(m0=0, v0=0, d1=-0.001, d2=0.001, S=-0.1, sigma=0.1, S2=-0.1))
getDataLikelihood(modelCoEvo_Split, simCoPM_geo,
                        c(m0=0, v0=0, d1=-0.001, d2=0.001, S=0, sigma=0.1, S2=-0.1))
~~~

These result in near identical log-likelihood values, showing that the JointPM_geo_ and CoEvo_split_ models collapse into the CoPM_geo_ model when *S_1_*=*S_2_*, and when *S_1_*=0, respectively.

We can further show that the CoEvo and CoEvo_all_ models are special cases of the CoEvo_split_ model.

~~~
simCoEvo <- simulateTipData(modelCoEvo_Split, method=1,
                c(m0=0, v0=0, d1=-0.001, d2=0.001, S=-0.1, sigma=0.1))
simCoEvo_all <- simulateTipData(modelCoEvo_all, method=1,
                c(m0=0, v0=0, d1=-0.001, d2=0.001, S=-0.1, sigma=0.1))
~~~

We then estimate the likelihood for CoEvo and CoEvo_split_ models to the first dataset, and CoEvo_all_ and CoEvo_split_ models to the second dataset.

~~~
getDataLikelihood(modelCoEvo, simCoEvo,
                        c(m0=0, v0=0, d1=-0.001, d2=0.001, S=-0.1, sigma=0.1))
getDataLikelihood(modelCoEvo_Split, simCoEvo,
                        c(m0=0, v0=0, d1=-0.001, d2=0.001, S=-0.1, sigma=0.1, S2=0))

getDataLikelihood(modelCoEvo_all, simCoEvo_all,
                        c(m0=0, v0=0, d1=-0.001, d2=0.001, S=-0.1, sigma=0.1))
getDataLikelihood(modelCoEvo_Split, simCoEvo_all,
                        c(m0=0, v0=0, d1=-0.001, d2=0.001, S=-0.1, sigma=0.1, S2=-0.1))
~~~

**Figure S1:**
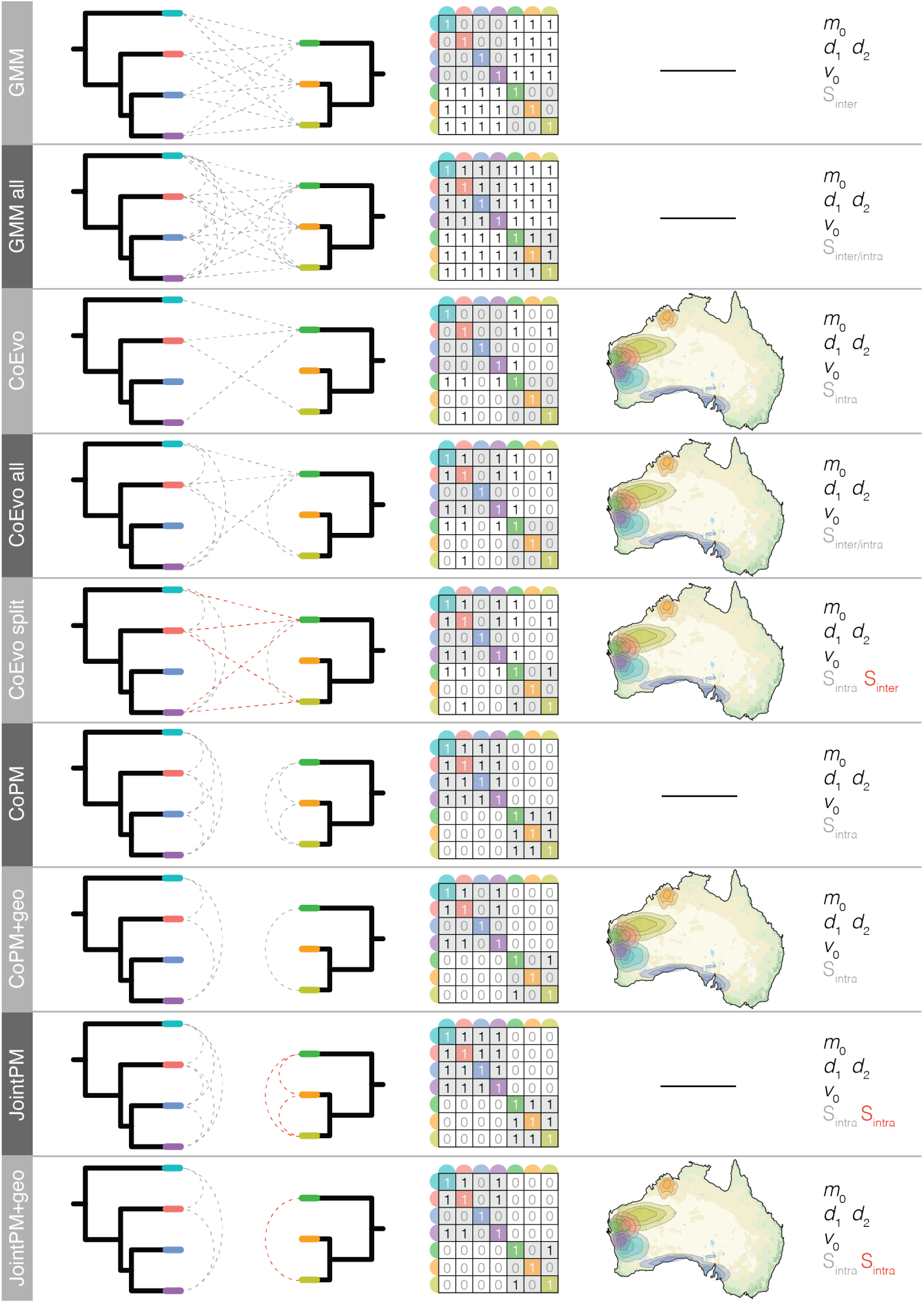
Schematic components of various GMM-type models of evolution including two clades. Each model name is listed at left, followed by a diagram of the two trees with interlineage interactions allowed under the given model designated by dashed grey lines. If more than one interaction parameter *S* is estimated, it is denoted by red dashed lines. The contemporary summary of these interactions are presented in the interaction matrix *P*, and the estimated parameters are listed at the far right. If the interaction matrix is geographically informed, a map showing species ranges is shown to the right of the interaction matrix.

**Figure S2:**
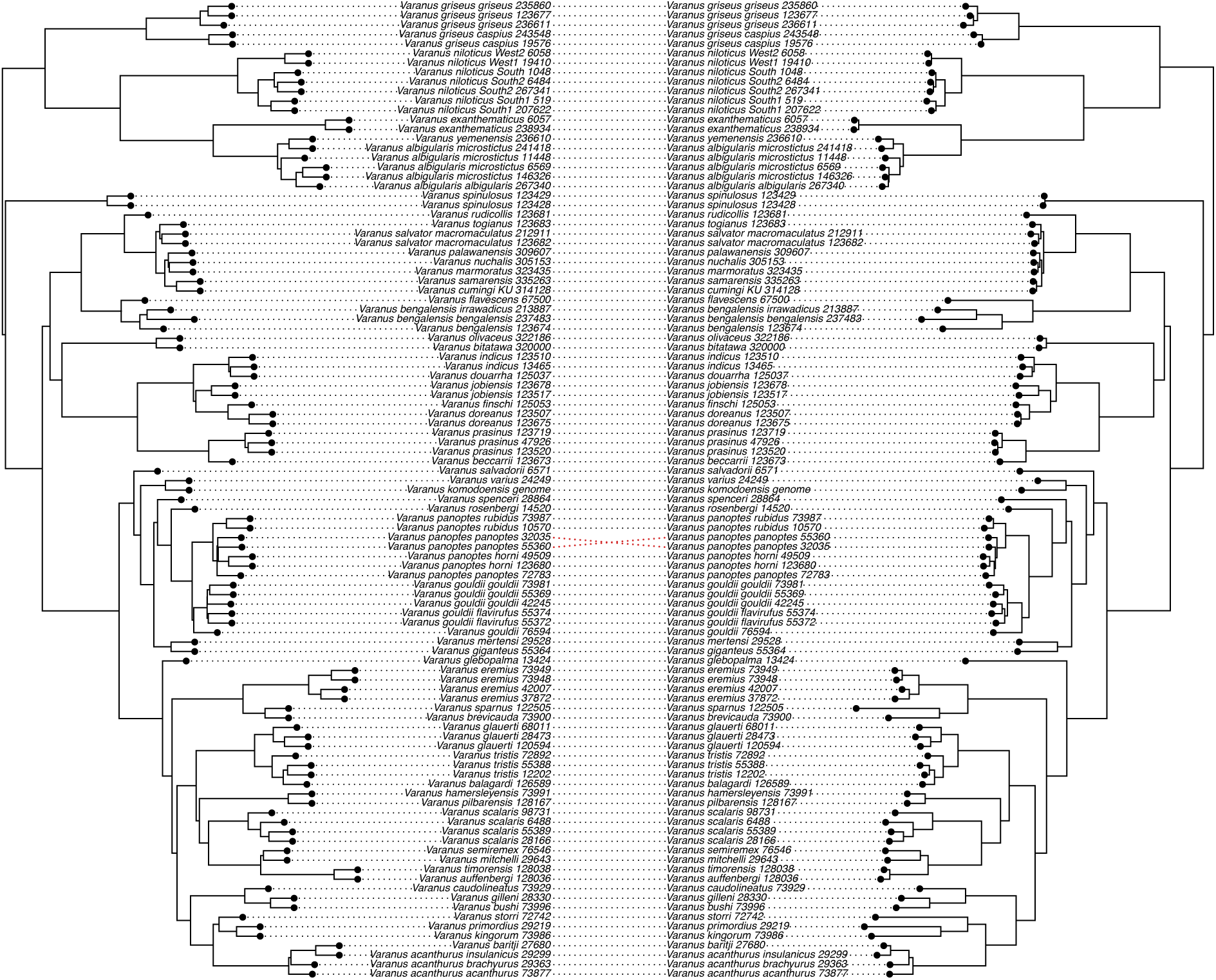
Phylogenetic estimates from shortcut coalescent (ASTRAL—left) Zhang et al. (2017) and partitioned maximum likelihood (IQTREE–right) Schmidt et al. (2014) analyses return nearly identical topologies.

**Figure S3:**
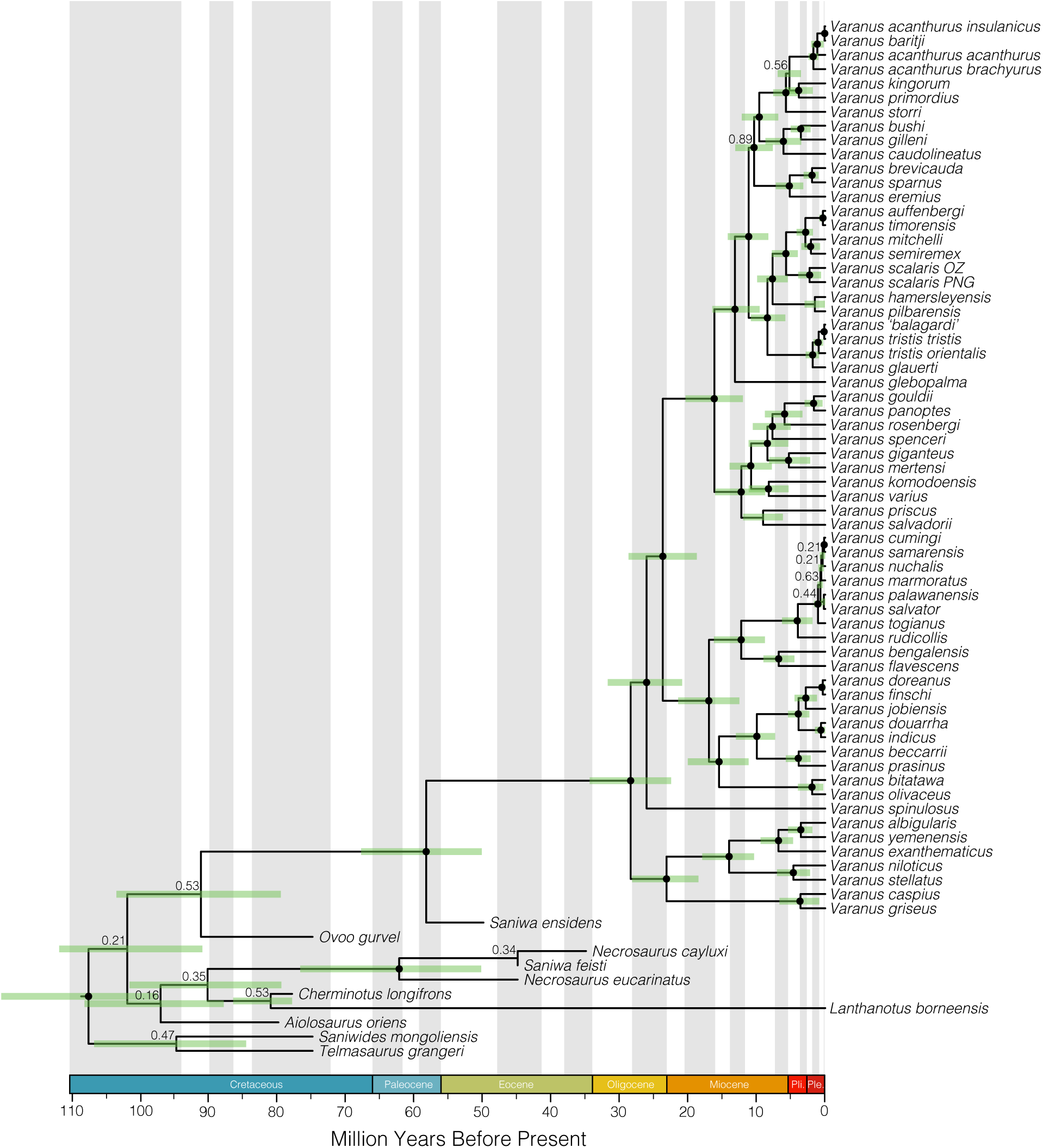
Relationships among living and extinct varaniform lizards and relatives, as a result of combined evidence dating (molecular and morphological data). The resolution of fossil taxa are volatile and appear highly sensitive to the fragmentary remains of many of these taxa. Varanids emerge in the late Paleocene or early Eocene, and extant *Varanus* appear in the Oligocene. Support values for relationships among *Varanus* subgenera as well as interspecific relationships are consistently high, though extinct *Varanus* are again difficult to place in our phylogeny. Nodes denoted by *•* are supported by posterior probabilities >0.90, all others are considered equivocal and labeled with posterior probabilites.

**Figure S4:**
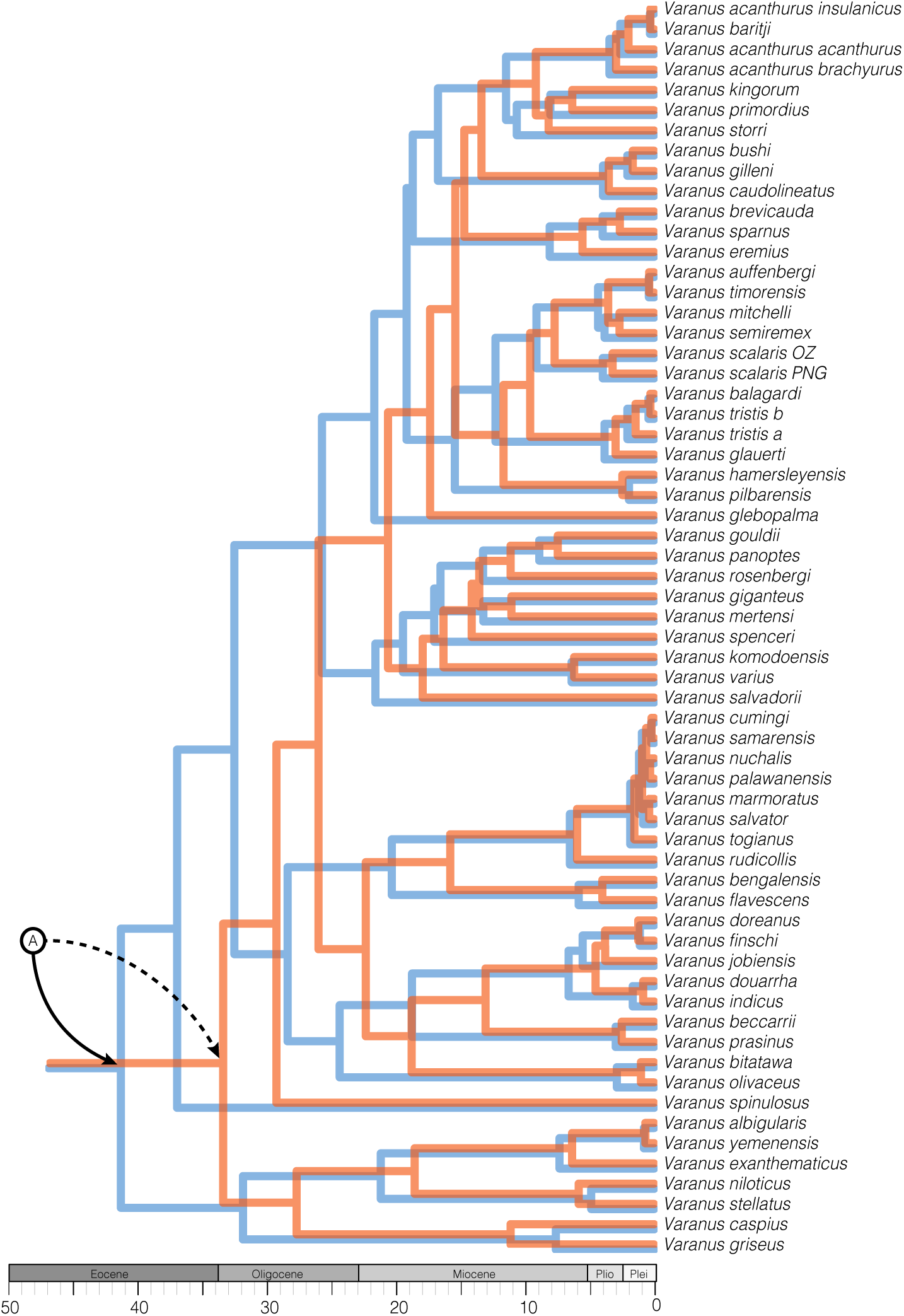
*V aranus* divergence estimates are strongly influenced by the inclusion of the ‘Jebel Qatrani *Varanus*’ (Holmes et al. 2010) as a minimum age node prior on crown *Varanus*. Morphological analyses have suggested that this taxon likely represents a stem varanid and incorporating this taxon as a node calibration prior inflates divergence times across the tree. The two presented trees are the result of calibrated multispecies coalescent analyses in StarBEAST2 using only extant taxa and molecular data. (A) denotes the position of the *Varanus* crown, the blue tree shows divergence times estimated using the ‘Jebel Qatrani’ fossil calibration, and the orange tree shows without. See further discussion of fossil taxa in *Node Priors and* Varanus *in the Fossil Record* in the Supplementary Material, and Table S8.

**Figure S5:**
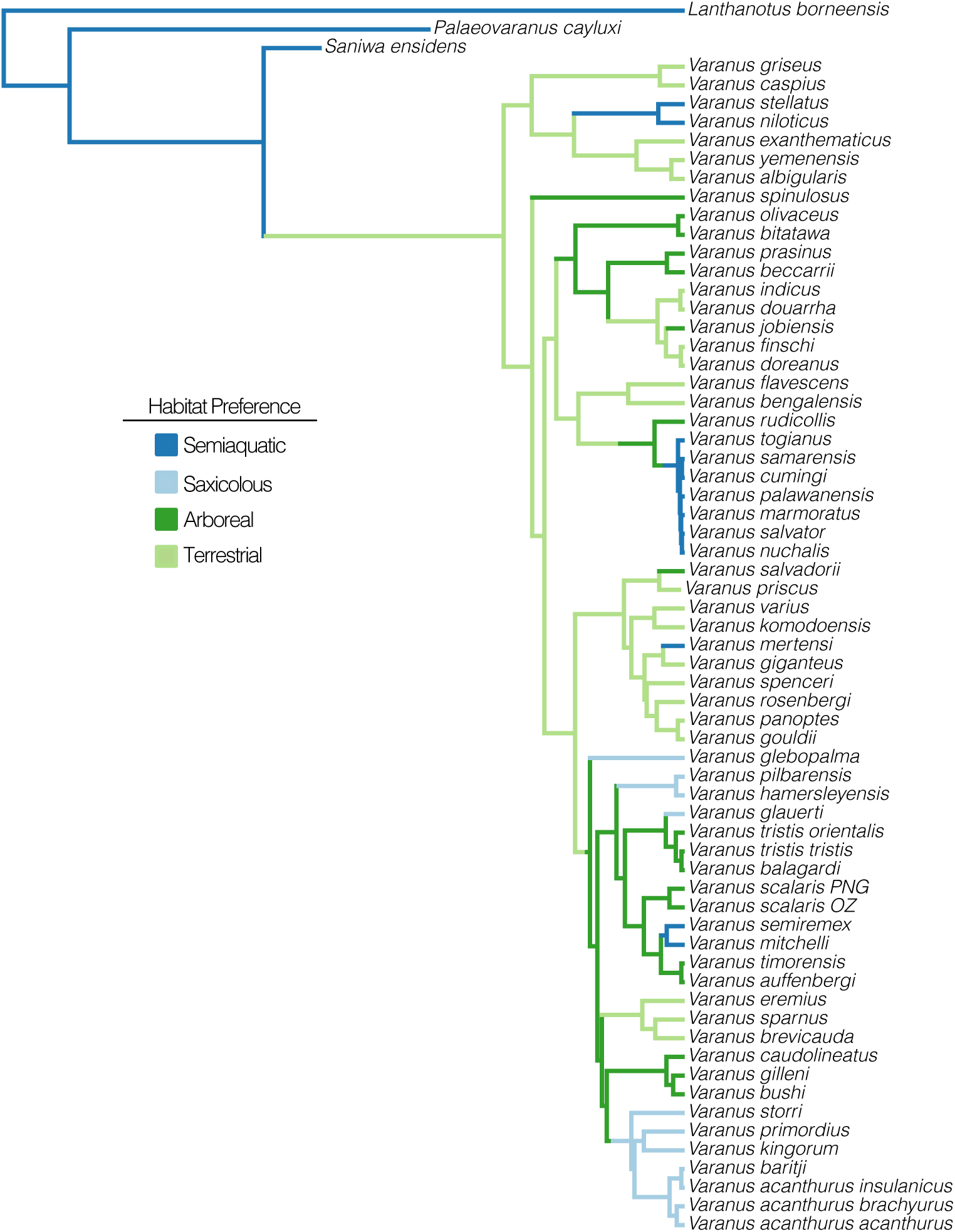
A single SIMMAP representation of the distribution of ecologies across *Varanus* lizards.

**Figure S6:**
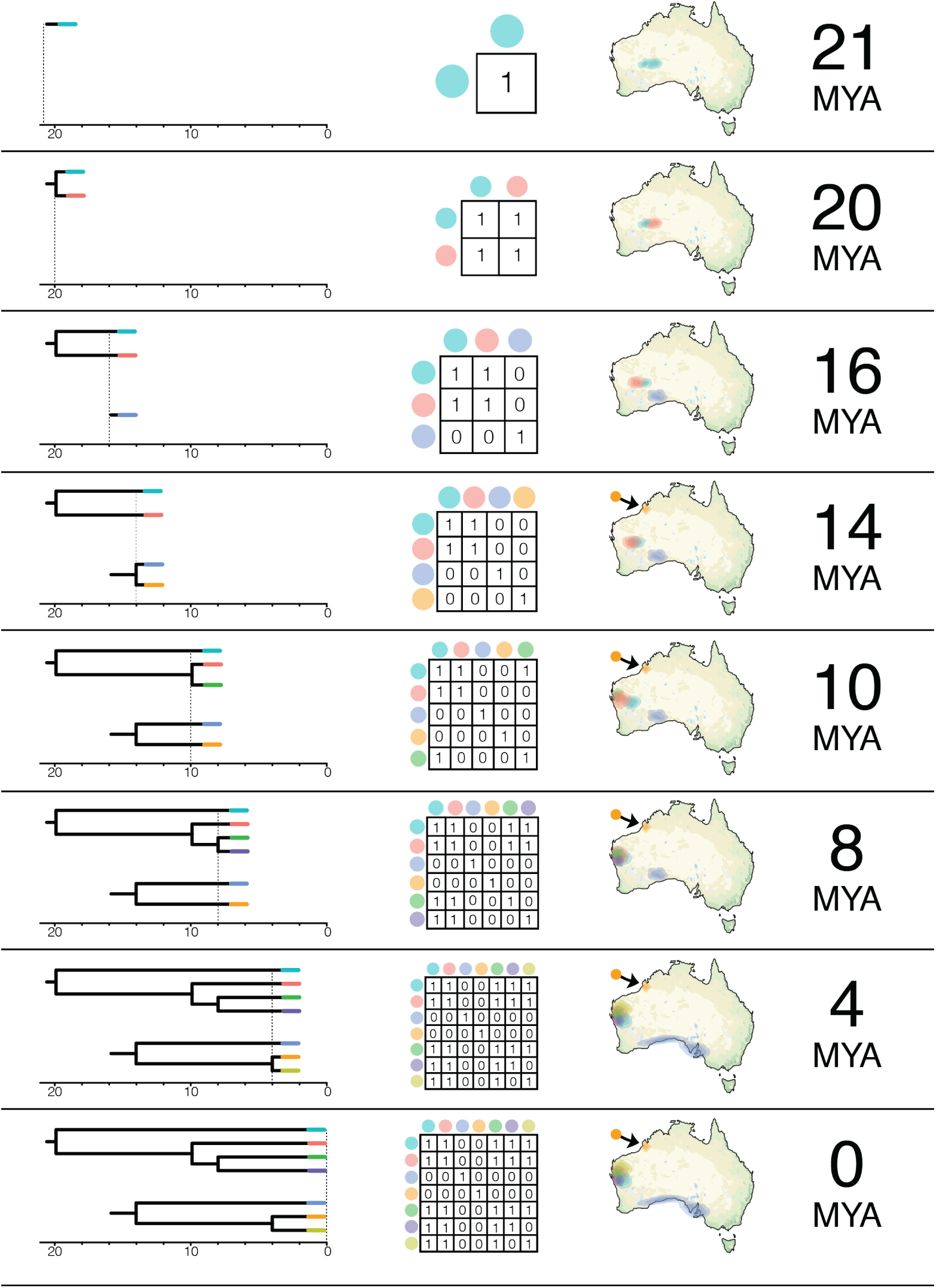
Diagram of the construction of interaction matrices *P* through time in geographically-informed models, using the CoEvo_all_ model as an example. Ancestral ranges were estimated using *rase*. The process of constructing these matrices is incorporated into the function “CreateCoEvoGeoObject_SP”, which takes as input the the trees, and two processed *rase* objects—one for each clade.

#### Investigating Data Completeness and Informativeness

Below we visualize data completeness and informativeness on a per sample and per locus basis, as well as provide some insight into our data cleaning and sample selection.

**Figure S7:**
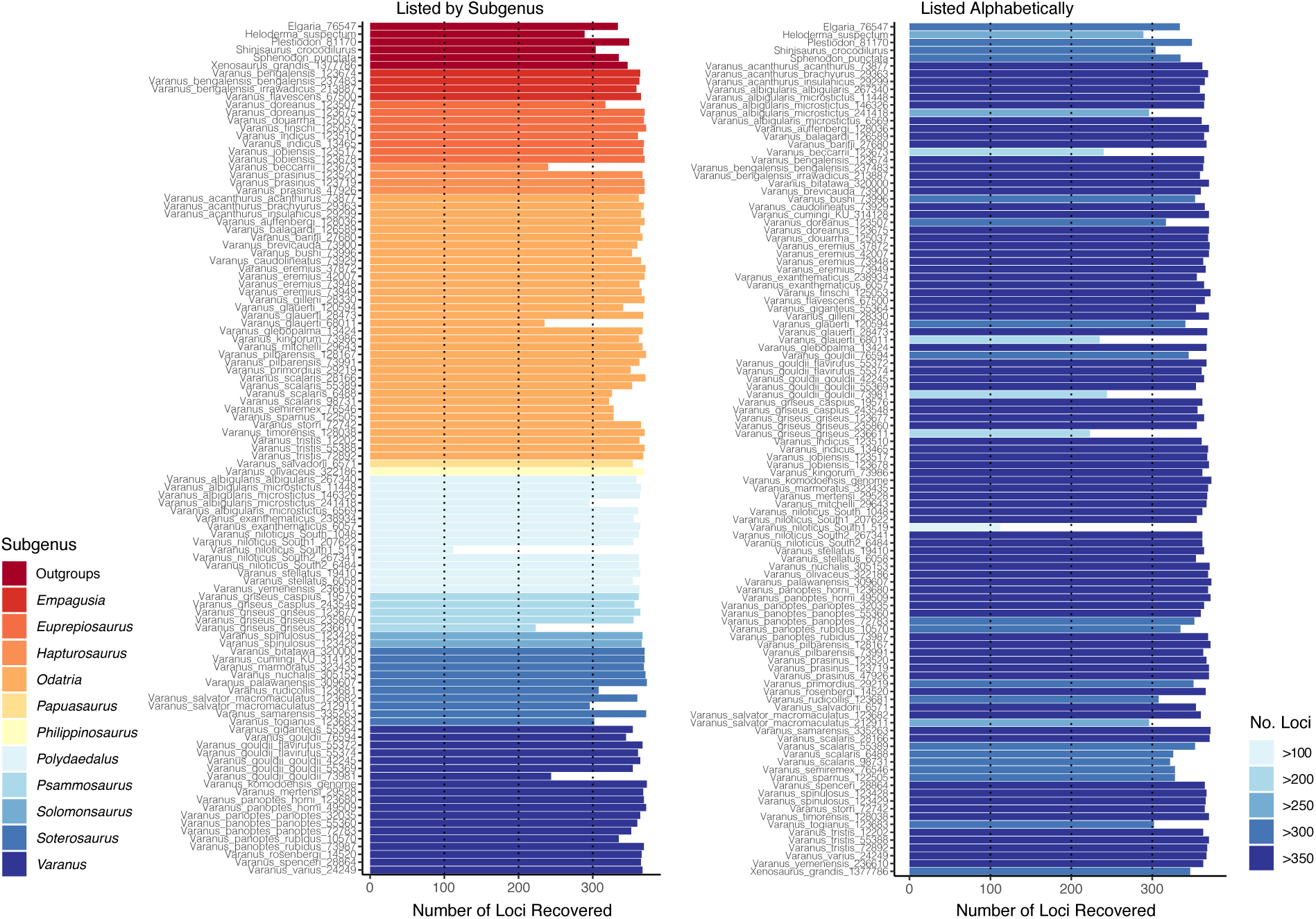
Number of loci recovered per sample for all *Varanus* and outgroup taxa included in the molecular data. Samples are ordered by subgenus (left) and alphabetically (right).

**Figure S8:**
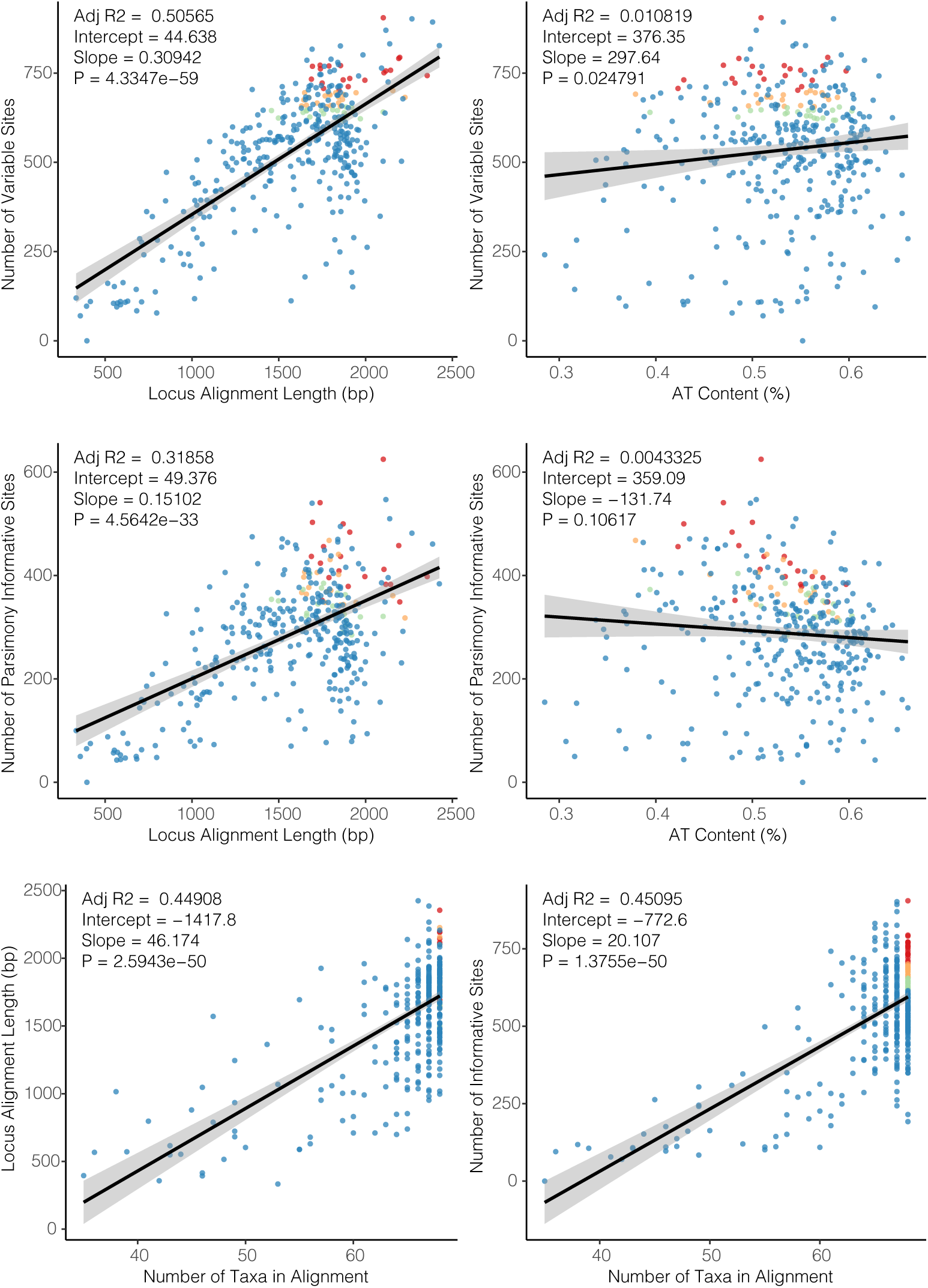
Plots of individual locus completeness and informativeness. For the StarBEAST2 species tree analyses, loci were ordered first by completeness (number of taxa in alignment), then by variable sites. They were then partitioned into 3 sets of twenty loci, and are color coded in these plots: ‘top twenty’ (1-20: red), ‘second twenty’ (21-40: orange), ‘third twenty’ (41-60: green), and all others (blue). Top row shows the number of variable sites in each alignment as a function of alignment length and AT content. The middle row shows the number of parsimony informative sites as a function of alignment length and AT content. The bottom row shows alignment length and number of variable sites as a function of completeness.

**Figure S9:**
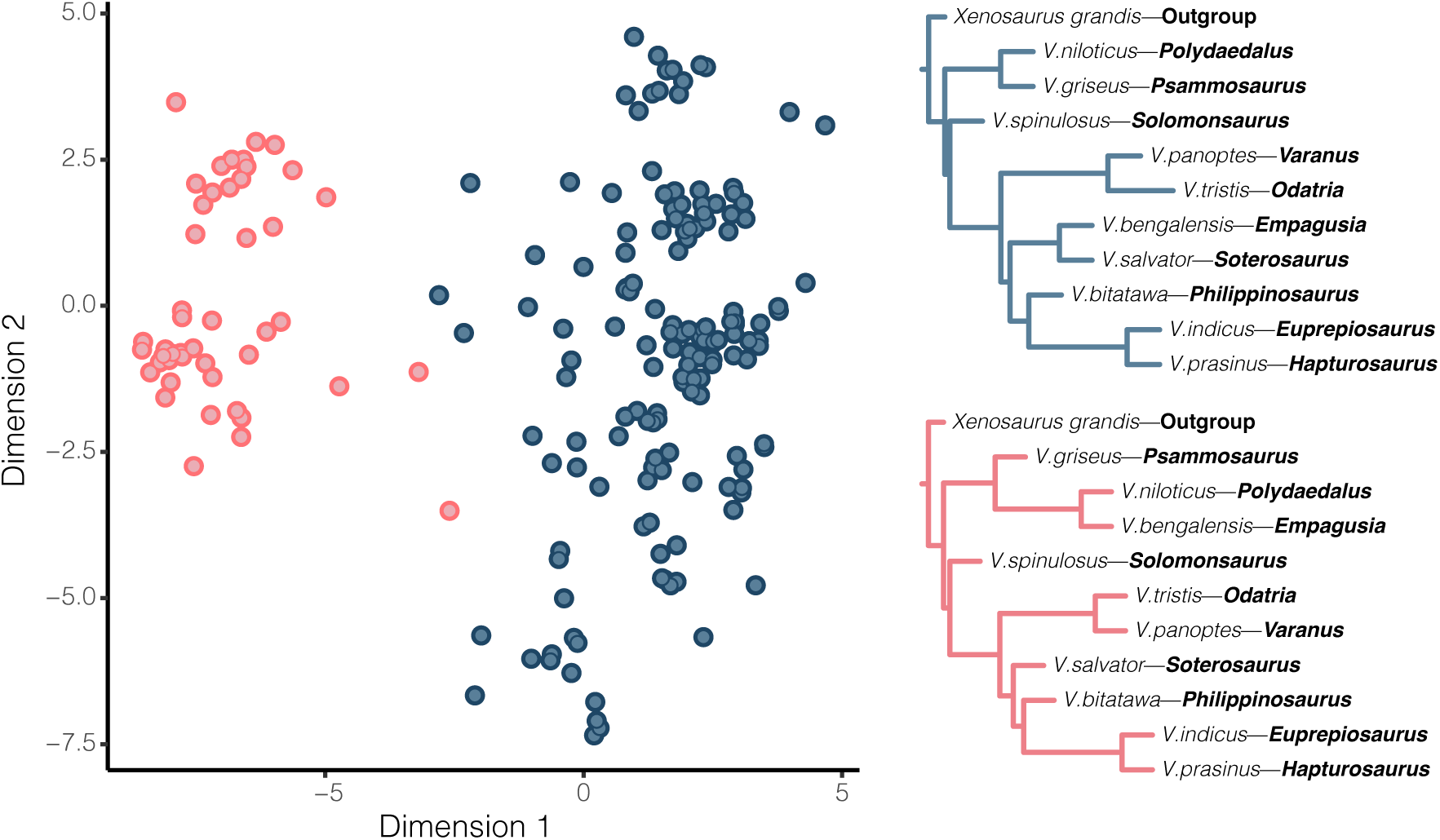
Two dimensional representation of multidimensional scaling (MDS) of gene tree space, colored by optimal clustering scheme (*k*=2), and their associated topologies inferred using ASTRAL. Analysis in both two and three dimensions supported the same optimum number of clusters, and cluster compositions. Each point represents a single gene tree, colored clusters match colored trees displayed to the right. Bootstrap support of all nodes was 1. The general topology of the clusters differ only in the placement of *V. bengalensis*—*Empagusia* as sister to the African group *Polydaedalus*, or to the Asian group *Soterosaurus*.

Preliminary analysis of genealogies indicated some strongly conflicting topologies between *Varanus* subgenera. To address gene-tree incongruence and investigate possible conflicting signals in our data, we used multidimensional scaling (MDS) to approximate the relative distances between gene tree topologies (Hillis et al. 2005), following the methodology of Duchene et al. (2018). To prepare the data, we trimmed down gene trees to a single representative of most subgenera (except *Papuasaurus*—*V. salvadorii*) as well as the outgroup *Xenosaurus*, and discarded loci missing any taxa, leaving us with 340 loci. We then calculated the pairwise distances between all gene trees using the Robinson-Foulds metric, in the R package APE (Paradis et al. 2004). We projected the tree distances into two and three dimensions (representing tree topology space) using MDS, as visualizing and interpreting any more dimensions becomes difficult. To test if gene trees are uniformly distributed throughout tree space, or clustered, we used the partitioning around medoids algorithm as implemented in the R package CLUSTER (Maechler et al. 2018). We chose the optimum number of clusters (*k*), using the gap statistic, calculated for each *k* = 1–10. Clusters of gene trees represent similar topologies, and so we then summarized each cluster using ASTRAL, to identify consistent differences in topology.

Multidimensional scaling (MDS) of gene-trees reveals that nuclear loci constitute two topological clusters. The larger cluster (*n*=264 loci) supports a sister relationship between *Empagusia* and *Soterosaurus*, and the smaller cluster (*n*=76 loci) supports a sister relationship between *Empagusia* and *Polydaedalus* (Fig.S9). Looking at fully-sampled gene trees we see that these patterns are driven by a sister relationship between *V. bengalensis* and *V. flavescens* (both *Empagusia*) in the larger cluster, and a sister relationship between *V. bengalensis* and *V. albigularis/V. yemenensis* in the smaller cluster.

This mixed-ancestry sample of *V. bengalensis* is perhaps interesting with regards to the large ranges of some African/Middle Eastern and mainland Asian *Varanus*. Previous research has highlighted the dispersal abilities of monitor lizards and shown that at least one member of the African varanids *Polydaedalus*—*V. yemenensis* has since dispersed back across the Red Sea into the Arabian Peninsula (Portik and Papenfuss 2012). On a similar time frame, *V.bengalensis* appears to have dispersed west back across Asia, and the subcontinent, into the Middle East. This is relevant because the mixed phylogenetic signature (Fig.S9) between one sample of *V. bengalensis* and members of the *V. albigularis* group, to which *V. yemenensis* belongs, suggests either introgression between these taxa, or a potentially contaminated sample. It remains an exciting concept that secondary contact between distantly related *Varanus* could result in hybridization, perhaps facilitated by the noted chromosomal conservatism of this genus (King and King 1975).

#### Testing for Fossil Taxa as Sampled Ancestors

Given that we place a prior on the age of each taxon (*τ*) and are jointly estimating their position among the phylogeny, including a model (*M*) of the molecular and morphological evolution, we can sample exclusively from both the prior and posterior of our starBEAST2 analyses. We used a threshold of log(BF) > 1 to identify sampled ancestors, log(BF) < −1 to recognize terminal taxa, and −1 < log(BF) < 1 taxa were categorized as equivocal. To calculate Bayes Factors for fossil taxa as sampled ancestors:

**Figure S10:**
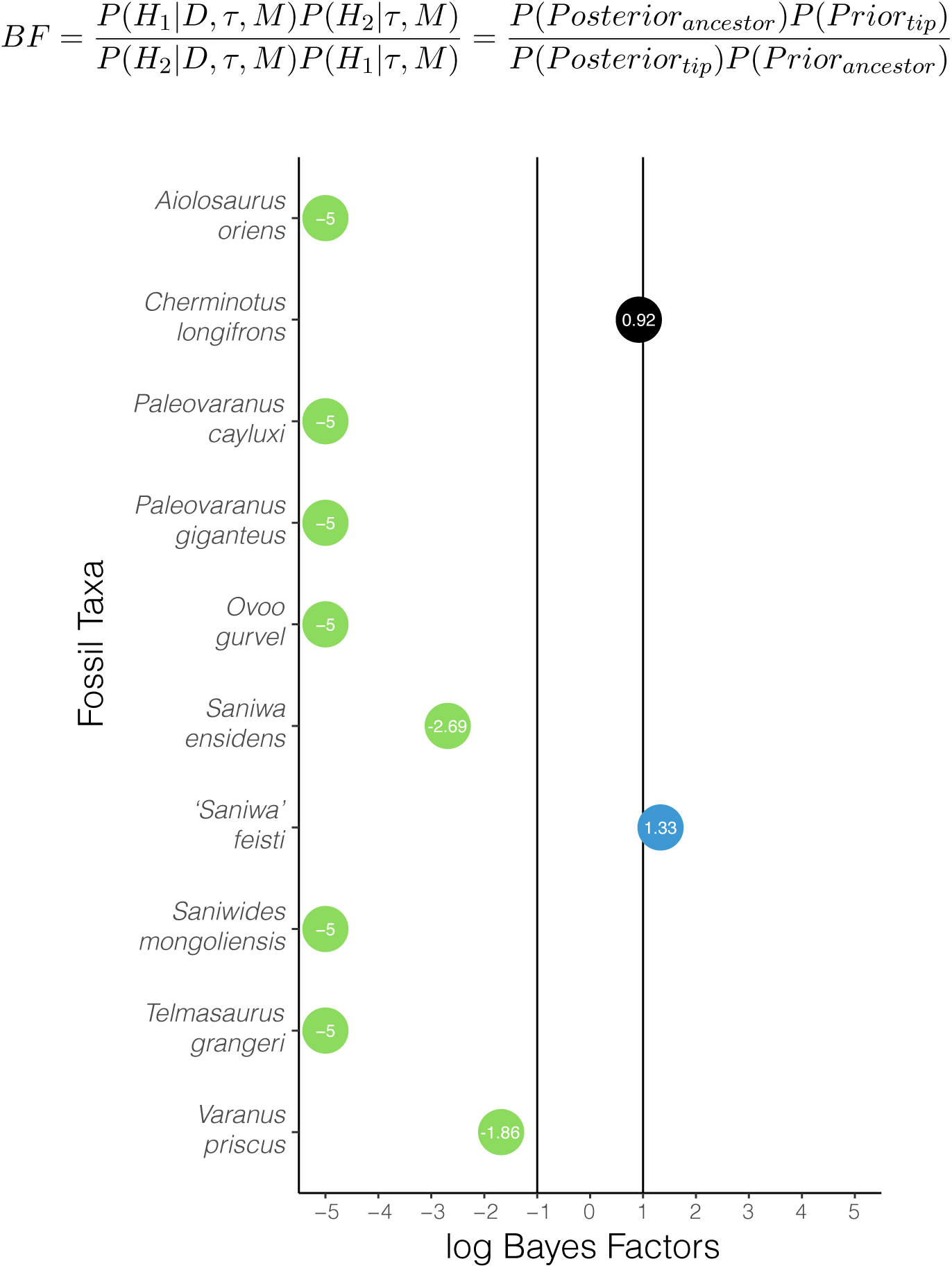
Bayes Factors support the position of nearly all fossil taxa as terminals. Green circles are strongly supported as terminal taxa, and black circles denote equivocal assignment. Very low log BF scores (taxa nearly always sampled as terminals) are reported arbitrarily as -5 to facilitate visualization.

**Figure S11:**
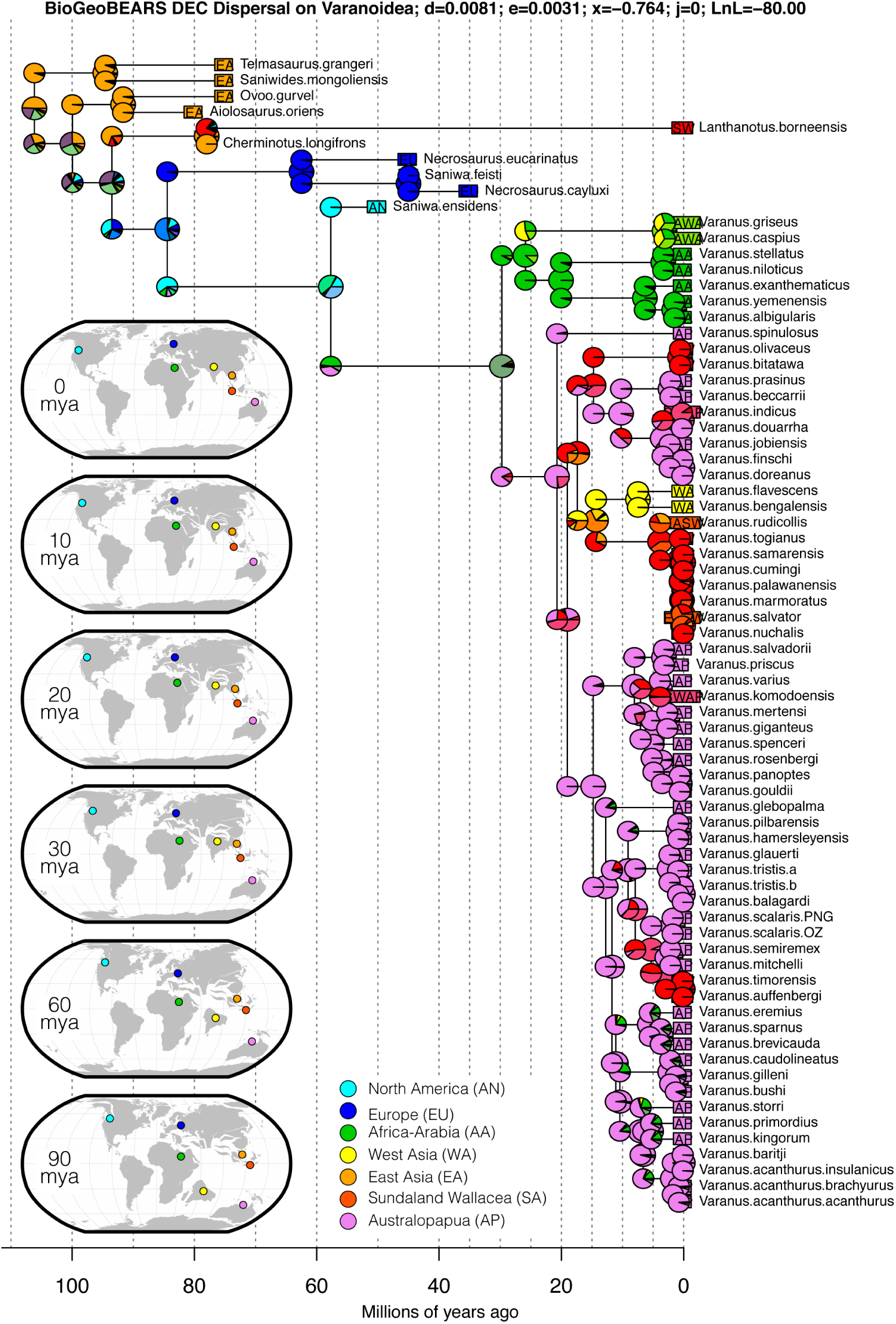
BioGeoBEARS ancestral state reconstruction under the DEC model with dispersal probability modelled as a function of distance among areas. Inset maps show the global position of major continents and land masses at relevant time slices (0, 10, 20, 30, 60, 90 million years ago). The ancestral distribution of varanoids is suggested to be Laurasian (East Asia + Europe), though members of this group are spread across all continents and subregions with the exception of South America and Antarctica. The ancestral distrbution of *Varanus* is highly ambiguous, and generally returns an estimate composed of all the major regions in which monitor lizards currently live. This is compounded by the enigmatic distribution of *Saniwa ensidens* in North America.

#### Model Fitting Results

**Figure S12:**
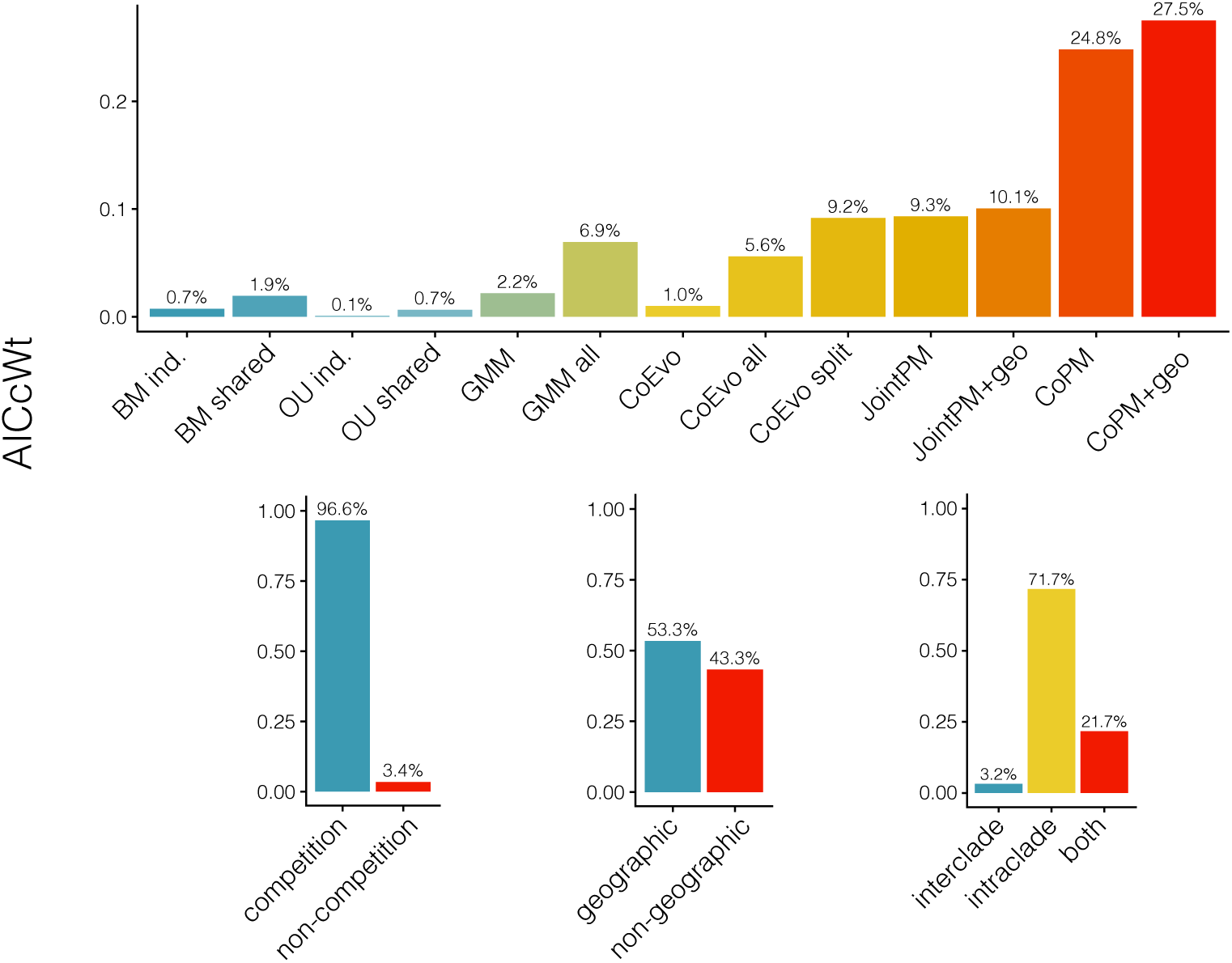
Comparative model fitting highlights the importance of incorporating interactions when modelling body size evolution of monitor lizards and dasyuromorphian marsupials. Modelling competition vastly improves model fit, but size evolution appears largely driven by intraclade evolution and not competition between monitors and mammals. Incorporating historical biogeography only narrowly improves model inference.

#### Model Identifiability

**Figure S13:**
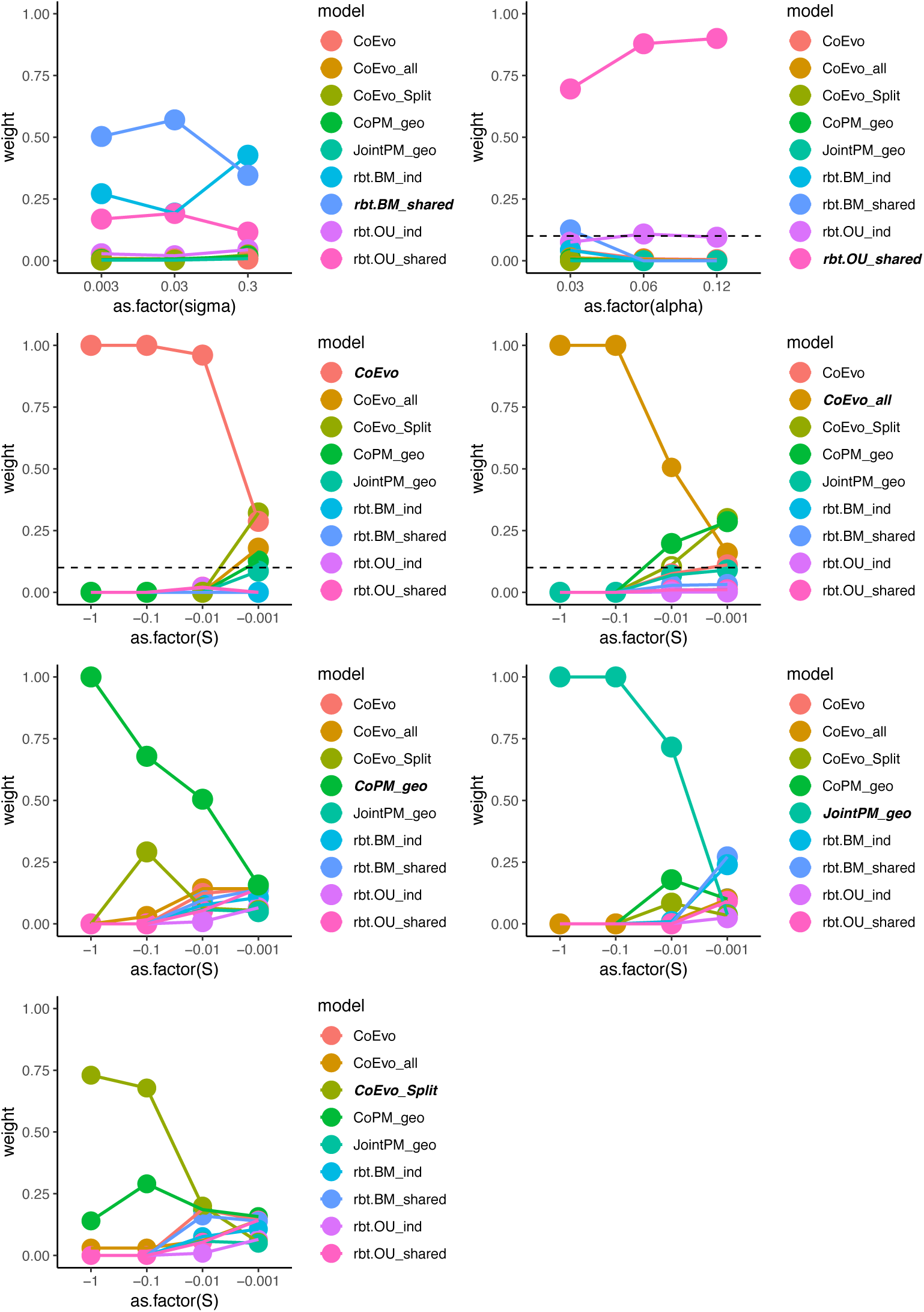
As the strength of competition *S* increases, model selection becomes more reliable. Results of model identifiability simulations as a function of varying parameter values. Identifiability (presented as AICCweight) of interaction models is uniformly poor for extremely small absolute values of *S*, but increases considerably at values of −0.01 and beyond. Values for simulations are included in Table S4.

#### Parameter Estimation Under GMM-type Models

**Figure S14:**
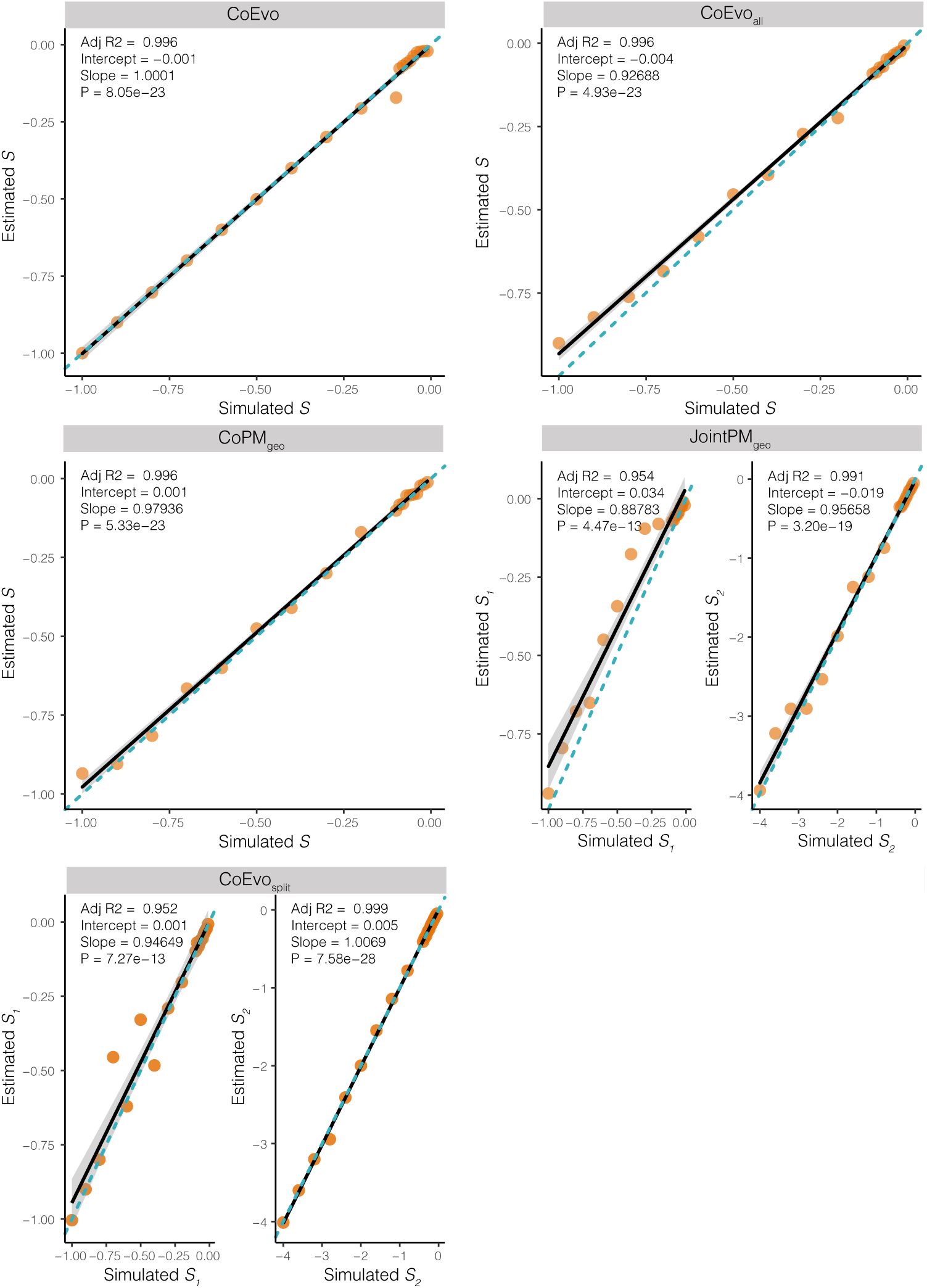
The competition parameter *S* can be accurately estimated under competitive models. Simulated values were −0.01, −0.02, −0.03, −0.04, −0.05, −0.06, −0.07, −0.08, −0.09, −0.1, −0.2, −0.3, −0.4, −0.5, −0.6, −0.7, −0.9, −1. Estimated values are consistently accurate between these limits.

## Acknowledgments

We would like to thank the Keogh and Moritz lab groups at ANU for discussion and comments throughout the development of this project. Thanks to Alex Skeels for a crash-course in methods of spatial data and functional diversity, and as always Zoe K.M. Reynolds for troubleshooting scripts. We appreciate the opportunity provided by the Society for Systematic Biology to present this research in the Ernst Mayr symposium at the 2018 Evolution meeting in Montpellier, France. A considerable thank you to the curators and staff of the many Australian and international museums and databases (Atlas of Living Australia, Australian Museum, Museum and Art Gallery of the Northern Territory, South Australian Museum, Australian Biological Tissue Collection, Queensland Museum, Western Australian Museum, Australian National Wildlife Council, University of Michigan Museum of Zoology, Bernice P.Bishop Museum, California Academy of Sciences, Museum of Vertebrate Zoology at Berkeley, Port Elizabeth Museum, Royal Ontario Museum, and University of Kansas Natural History Museum) for access to tissues and locality data that made this work possible.

